# Chemoproteomic Profiling of *C. albicans* for Characterization of Anti-fungal Kinase Inhibitors

**DOI:** 10.1101/2025.01.10.632200

**Authors:** David J. Shirley, Meganathan Nandakumar, Aurora Cabrera, Bonnie Yiu, Emily Puumala, Zhongle Liu, Nicole Robbins, Luke Whitesell, Jeffrey L. Smith, Scott P. Lyons, Angie L. Mordant, Laura E. Herring, Lee M. Graves, Rafael M. Couñago, David H. Drewry, Leah E. Cowen, Timothy M. Willson

**Affiliations:** Structural Genomics Consortium and Division of Chemical Biology and Medicinal Chemistry, UNC Eshelman School of Pharmacy, University of North Carolina at Chapel Hill, Chapel Hill, NC 27599, USA; Department of Pharmacology, University of North Carolina at Chapel Hill, Chapel Hill, NC 27599, USA; UNC Lineberger Comprehensive Cancer Center, School of Medicine, University of North Carolina at Chapel Hill, Chapel Hill, NC 27599, USA; Department of Molecular Genetics, University of Toronto, Toronto, ON, M5G 1M1, Canada; Center of Medicinal Chemistry, Center for Molecular Biology and Genetic Engineering, University of Campinas, 13083-886-Campinas, SP, Brazil

**Keywords:** multiplexed inhibitor beads (MIBS), multiplexed inhibitor beads analyzed by tandem mass-spectrometry (MIB/MS), chemoproteomics, antifungal, kinase inhibitor, in-cell target engagement assay

## Abstract

*Candida albicans* is a growing health concern as the leading causal agent of systemic candidiasis, a life-threatening fungal infection with a mortality rate of ∼40% despite best available therapy. Yck2, a fungal casein kinase 1 (CK1) family member, is the cellular target of inhibitors YK-I-02 (YK) and MN-I-157 (MN). Here, multiplexed inhibitor beads paired with mass spectrometry (MIB/MS) employing ATP-competitive kinase inhibitors were used to define the selectivity of these Yck2 inhibitors across the global *C. albicans* proteome. The MIB matrix captured 89% of the known and predicted *C. albicans* protein kinases present in cell lysate. In MIB/MS competition assays, YK and MN demonstrated exquisite selectivity across the *C. albicans* fungal kinome with target engagement of only three CK1 homologs (Yck2, Yck22, and Hrr25) and a homolog of human p38α (Hog1). Additional chemoproteomics using a custom MN-kinobead identified only one additional *C. albicans* protein, confirming its remarkable fungal proteome-wide selectivity. To identify new Yck2 inhibitors with selectivity over Hog1, thirteen human CK1 kinase inhibitors were profiled for fungal kinase-binding activity using MIB/MS competition assays and in-cell NanoBRET target engagement assays. A new chemotype of family-selective Yck2 inhibitors with antifungal activity was identified. Together, these findings expand the application of MIB/MS proteomic profiling for non-human kinomes and demonstrate its utility in the discovery and development of selective inhibitors of fungal kinases with potential antimicrobial activity.

**GRAPHICAL ABSTRACT:** 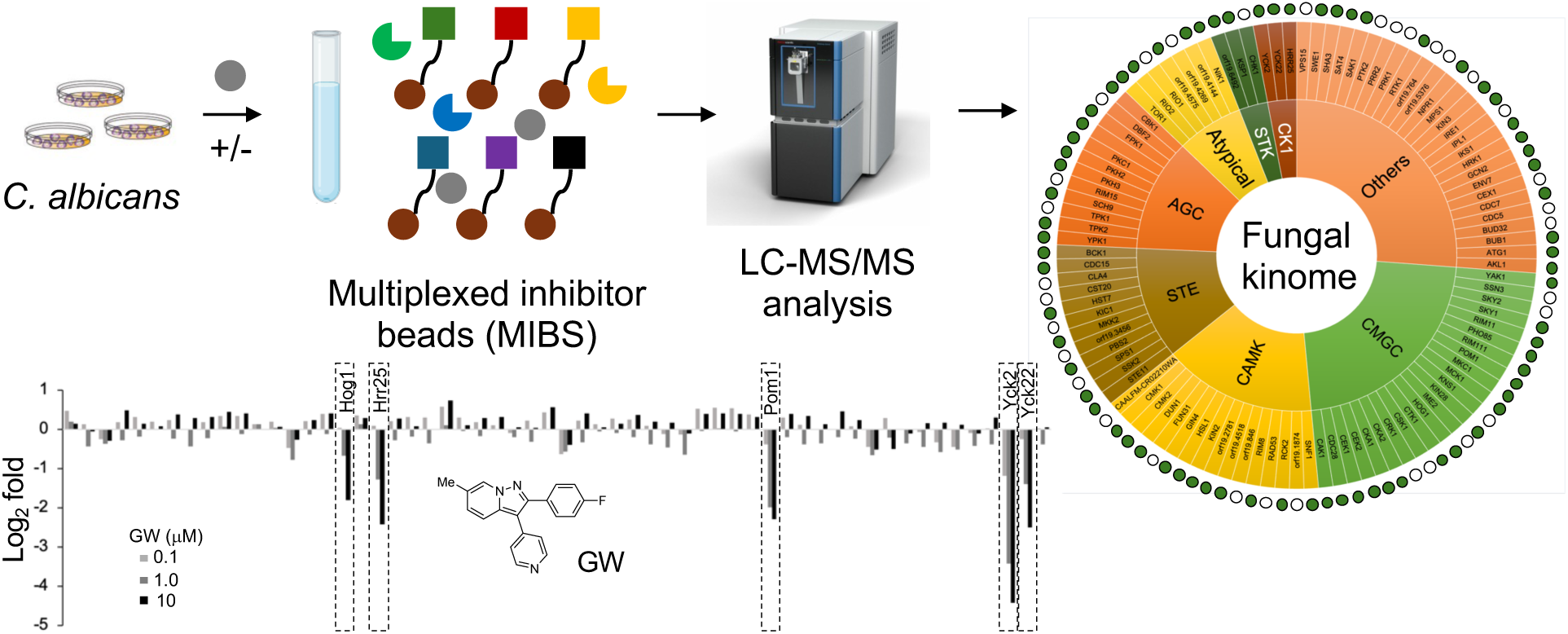

## INTRODUCTION

Pathogenic fungi have emerged as a global health concern, causing billions of infections and 2.5 million annual deaths^1,2^. Although over 600 different fungal species have been reported to infect humans, the W.H.O. has placed four species in the “Critical Priority Group”: *Cryptococcus neoformans*, *Aspergillus fumigatus*, *Candida auris*, and *Candida albicans*. Notably, *Candida* infections account for nearly 70% of all invasive fungal infection (IFI)-related deaths^3,4^. *C. albicans* is the most common causative agent of *Candida* infections with mortality rates ranging from 20–50% despite antifungal therapy^4^. IFI’s tend to be opportunistic, taking place in hospitalized patients as a complication of surgical procedures, as well as in patients with impaired immune function because of treatment with chemotherapeutics, HIV infection, or other immunocompromising disease^5^. The CDC estimates that these drug-resistant microbial infections cost the U.S. healthcare system approximately $4.6 billion annually^6^. Currently there are only four classes of antifungal drugs available to treat systemic infections: azoles, polyenes, flucytosine, and echinocandins^7–9^, with the emergence of drug resistance becoming a major clinical issue^7,10^. Yet, there has been limited investment in the discovery and development of new therapies due to concerns about commercial viability^3,11^. The identification and optimization of antifungal drugs acting by new molecular mechanisms has been further complicated by the close phylogenetic relationship between humans and fungi, which can make selective targeting of fungal proteins difficult. These compounding issues have led to an increase in *Candida* infections resistant to current drugs and an increased toll on the healthcare system. There remains an urgent need for discovery and development of new, targeted antifungal therapeutics.

Protein kinases are a target class known to be highly tractable for the development of orally active small molecule inhibitors. There are >80 FDA-approved kinase inhibitor drugs^12–14^. Fungal protein kinases play pivotal roles in signaling pathways that regulate growth, response to external stressors, morphogenesis, and antifungal drug-resistance, making them promising drug targets^15,16^. Remarkably, there has been no prospective targeting of the fungal protein kinome as an approach to developing new anti-infective drugs. Recently, our laboratory conducted a phenotypic screen of the published kinase inhibitor sets 1 & 2 (PKIS/ PKIS2) against *C. albicans* to uncover GW461484A (GW) as an antifungal agent that enhanced the efficacy of echinocandins against drug-resistant clinical isolates^17–19^. Haploinsufficiency profiling identified the primary molecular target as Yck2, a fungal homolog of casein kinase 1 (CK1), which regulates transcription factors involved in fungal morphogenesis, biofilm formation, host cell damage, and cell wall integrity^20^. Optimization of GW through structure-guided design and metabolic profiling produced lead compounds YK-I-02 (YK) and MN-I-157 (MN) with decreased hepatic clearance and improved pharmacokinetic properties^21^. YK and MN rescued HepG2 cell viability after *C. albicans* co-infection in culture and demonstrated antifungal activity in an immunocompromised mouse model of systemic *Candida* infection, both alone and in combination with the conventional antifungal caspofungin. However, further progression of these lead compounds as potential drug candidates would benefit from a better understanding of their mode of action, especially their selectivity across the fungal kinome.

Herein, we report the first use of multiplexed inhibitor beads (a.k.a. kinobeads) quantified by LC-MALDI TOF/TOF mass spectrometry (MIB/MS) for proteomic profiling of the *C. albicans* kinome. A MIB matrix utilizing six promiscuous ATP-competitive kinase inhibitors captured 89% of the predicted fungal protein kinases present in *C. albicans* and revealed the exceptional kinome-wide selectivity of YK and MN, with engagement of only three fungal CK1 homologs (Yck2, Yck22, and Hrr25) and a fungal p38 homolog (Hog1). Building on the phylogenetic similarity of eukaryotic kinomes, a diverse collection of human CK1 inhibitors were screened for *C. albicans* Yck2 inhibition, resulting in the identification of new chemotypes with selectivity for Yck2 over Hog1.

## RESULTS AND DISCUSSION

MIB/MS has emerged as a powerful chemoproteomic tool for analyzing inhibitor selectivity across the human kinome by measuring target engagement across hundreds of kinases expressed in cell lysates utilizing liquid chromatography coupled to tandem mass spectrometry (LC-MS/MS)^22–27^. MIB/MS competition assays utilize unmodified test compounds and require no *a priori* knowledge of their kinase target(s). A MIB matrix was previously developed that incorporates six promiscuous ATP-competitive inhibitors (Shokat, PP58, Purvalanol B, UNC-21474, VI-16832, and CTx-0294885) and routinely captures 75–80% of the expressed kinome in human cell lysates^28–30^. For our initial investigation we sought to define the fraction of the fungal kinome that was captured using MIB/MS by comparing results of this assay with a global analysis of the *C. albicans* proteome^31^. The global proteomic analysis of *C. albicans* lysate was performed by LC-MS/MS with the thermos Neo-Orbital Astral platform ^32–34^. The data was searched against the *C. albicans* strain SC5314 database from UniProtKB, which contains 6,039 open reading frames annotated as unique fungal proteins (Figure 1A)^35,36^. A total of 3,928 *Candida* proteins were identified in the cell lysate by this global proteomic analysis (Figure 1A, File S1).

**Figure 1.**
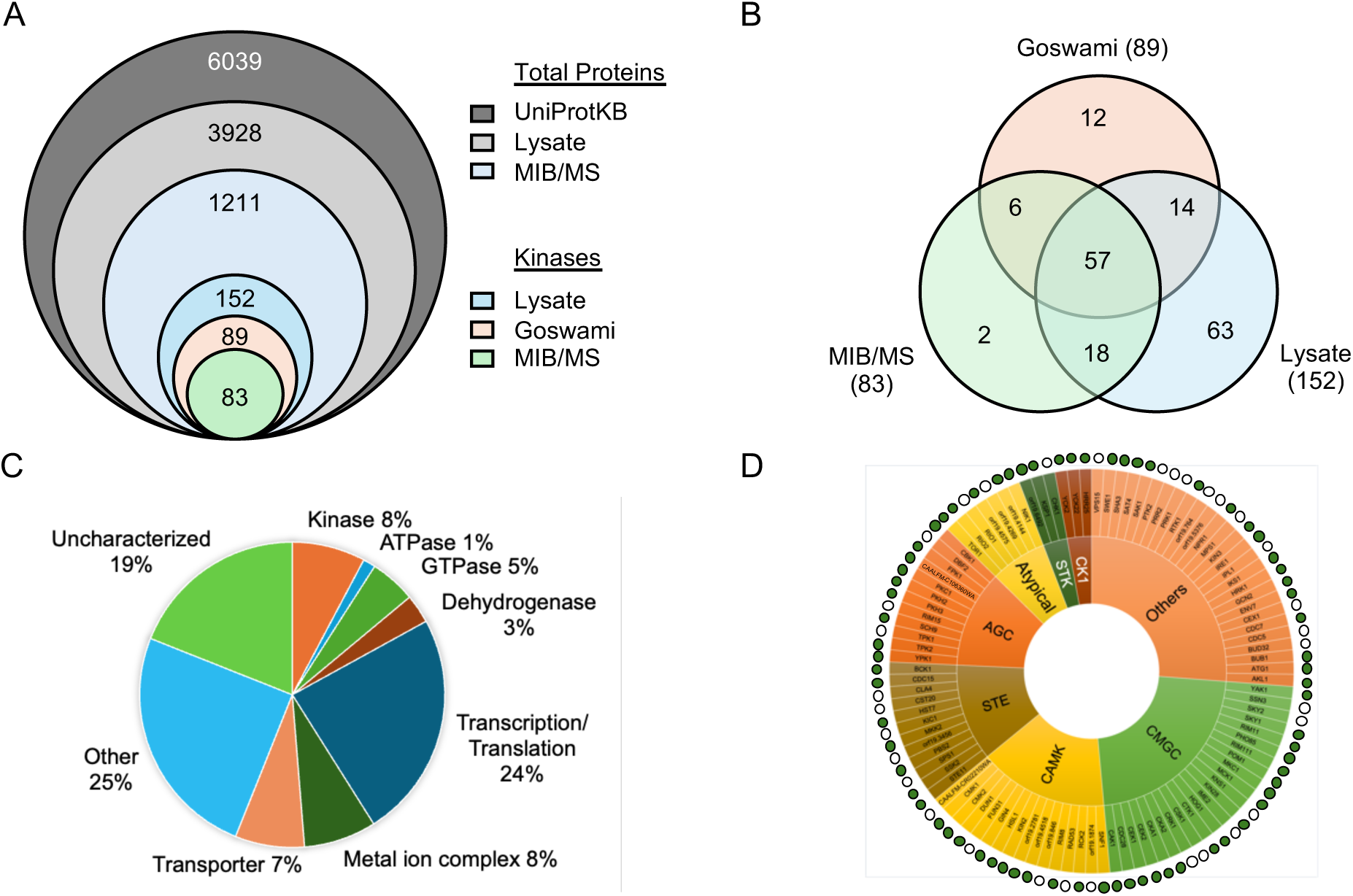
MIB/MS captures the majority of protein kinases in *C. albicans*. **A.** 6,039 proteins are predicted by UniProtKB to be encoded in the genome of *C.albicans*. 3,928 proteins were detected by global proteomic analysis. 1,211 of these proteins were captured by the MIB matrix of which 152 were annotated a potential kinase. 89 of these potential kinases were identified by Goswami et al., while 83 were captured by MIB/MS. **B.** Venn diagram displaying overlap between the Goswami annotation, MIB/MS experiments, and global proteomic analysis. **C.** Functional classification of the 1,211 fungal proteins captured by MIB/MS. **D.** The fungal protein kinome displayed by family. Kinases captured by MIB are shown by green dots on the sunburst plot. Open circles indicate kinases not captured by MIB.

The total size of the fungal kinome remains a subject of debate. UniProtKB lists 166 putative kinase-like proteins defined either by sequence similarity and computational annotation using ProtNLM or through experimental validation (e.g. mass spectrometry, NMR, X-ray determination, or Edman sequencing). The open-source Candida Genome Database lists 122 potential protein kinases using gene product readout, gene sequence, and direct reference to experimental findings^37^. Most importantly, Goswami and coworkers performed a rigorous analysis of the fungal kinome of several *Candida* species using Kinnanote software to identify 103 high confidence protein kinases when including both unusual and atypical kinases^38^. However, only 89 of the Goswami high confidence kinases are listed in UniProtKB, with the sequence of the other 14 not currently present in the database of predicted proteins. Our global proteomic analysis of *Candida* cell lysate identified 152 putative kinases through database matching of tryptic peptides using Spectronaut or MaxQuant software, of which a total of 71 are high confidence protein kinases as defined by Goswami (Figure 1C, File S1 and S2). The other 81 sequences were mostly non-protein metabolic kinases that phosphorylate sugars and nucleotides or proteins with low sequence homology that are unlikely to be functional protein kinases.

Next, the breadth of the fungal kinome captured by MIB was determined. After LC-MS/MS analysis, a total of 1211 fungal proteins were identified from the MIB/MS experiment using *C. albicans* cell lysate (File S2 and S3). Within this dataset, 83 potential *Candida* protein kinases were identified, of which 63 were high confidence protein kinases (Figure 1C). Six of these high confidence protein kinases were not identified in the global proteomic analysis, suggesting that they were expressed at low levels in the *C. albicans* lysate and had been enriched by the MIB matrix. Label-free quantification showed that protein kinases were enriched by ∼13-fold over other *Candida* proteins by the MIB matrix (File S2) In total, the MIB matrix captured 63 of 71 high-confidence protein kinases (89%) present in the *Candida* cell lysate (Figure 1C). Importantly, the MIB matrix captured a majority of kinases within each of the major families (Figure 1D) further demonstrating the ability of bead-immobilized ATP-competitive inhibitors to engage the full diversity of the *Candida* kinome. Notably, many other nucleotide binding proteins were also captured by the MIB matrix, including 16 ATPases, 59 GTPases, and 37 dehydrogenases. In addition, 291 proteins involved in transcription and translation (e.g. helicases, polymerases, ligases, and ribosome subunits) as well as several heme-containing proteins were captured (Figure 1B). Homologous ATP-binding proteins have often been captured from human cell lysates^39,40^. These findings were reproducible across subsequent experiments with 60 high-confidence protein kinases routinely identified by MIB/MS from fungal cell lysate.

Having defined the *C. albicans* kinome captured by MIB/MS, a series of competition experiments were performed with three Yck2 inhibitors (GW, YK, and MN) that have demonstrated robust antifungal activity^21^. In each experiment the MIB matrix captured 60–63 high confidence protein kinases from the fungal cell lysate for analysis. The antifungal compounds were remarkably selective across the fungal kinome (Figure 2), with a concentration-dependent competition by inhibitors observed for only a handful of protein kinases. GW, YK, and MN each competed for binding of binding of Yck2, Yck22, Hrr25, and Hog1 to the MIB matrix. In addition, GW competed with Pom1. Yck2, the previously identified fungal kinase target, is a CK1 homolog responsible for regulating *C. albicans* biofilm formation, host-cell damage, morphogenesis, and cell wall integrity^17,20^. Yck22 is the closest fungal paralog to Yck2. Although its function remains undefined, some studies have suggested that its activity is partially redundant with Yck2^41,42^. Hrr25 is a third fungal CK1 paralog and has been shown to regulate responses to both DNA and cell wall damage^43^. Hog1 is the fungal homolog of the human p38 MAPK, recognized for regulating cellular responses to oxidative and osmotic stress^44^. Notably, GW was originally designed as a human p38α kinase inhibitor^45^. GW, YK, and MN retained this MAPK activity in the fungal kinome through their engagement of Hog1. *C. albicans* Pom1, a homolog of the human dual-specificity tyrosine-regulated kinase (DYRK), has been shown to be important for the yeast-to-filament transition, a development switch closely linked to virulence^46,47^. YK and MN did not engage Pom1 and, thus, have improved fungal kinome-wide selectivity compared to GW. Although all three fungal kinase inhibitors showed binding to <10 non-kinase, nucleotide binding proteins, none of these interactions were observed with more than one of the inhibitors (File S4a and b), further underscoring the remarkable selectivity of the YK and MN chemotypes for only the Yck2 paralogs and Hog1 protein kinases.

**Figure 2.**
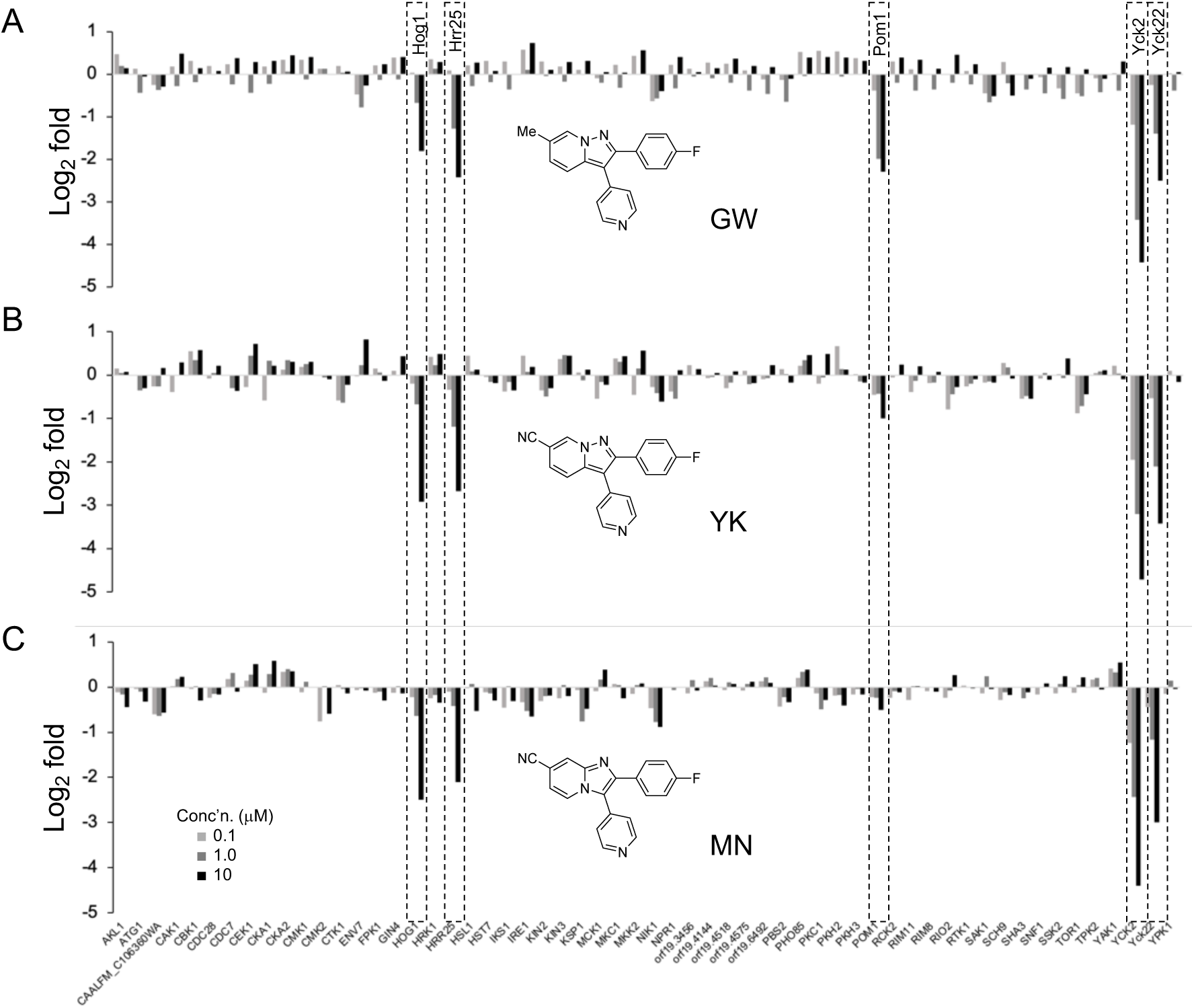
GW, YK, and MN demonstrate high selectivity across the fungal kinome. Fold change of fungal protein kinase targets captured in MIB/MS competition assays for each compound compared to DMSO. Assays were performed in triplicate.

Because MIB profiling of the fungal cell lysates was performed using a bead matrix that employed only six promiscuous kinase inhibitors, it remained possible that there were additional fungal kinase targets of GW, YK, and MN that were not detected in the proteomic analysis. Specifically, there were 14 protein kinases identified by Goswami that were present in *C. albicans* lysate but were not enriched by the MIB matrix and an additional 12 that were not captured in the global proteomic analysis (Figure 1C). Therefore, a reverse proteomic experiment was designed using an ECH-Sepharose 4B bead with MN covalently attached to explore whether there might be additional fungal-specific proteins that interact with the inhibitor. Structure-activity of the MN chemotype had established that modifications at the 6- or 7-position of the imidazo[1,2-*a*]pyridine retained antifungal activity^21^. A MN-loaded Sepharose bead (MN-kinobead) was synthesized from 7-methoxy-MN^21^ (Figure 3A), washed, and stored in a 20% ethanol solution. To demonstrate the fidelity of the MN-kinobead, a pull down was performed using human cell (HEK293) lysates. The MN-kinobeads mirrored the ability of positive control PP58 kinobeads^45^ to capture p38α as shown by Western blotting whereas bare Sepharose beads failed to pulldown the kinase (Figure 3B and S1). Incubation of the fungal lysate with the MN-kinobeads followed by competition with free MN (10 µM) demonstrated target competitive binding of only four protein kinases Yck2, Yck22, Hrr25, and Hog1, mirroring the results obtained with the MIB matrix (Figure 3C, File S5). The only non-kinase fungal protein that demonstrated specific binding to the MN-kinobead was CAALFM_C401020, a putative PIN domain RNA nuclease involved in nonsense mediated mRNA decay^48,49^. These results further highlight the exquisite selectivity of MN for the three *Candida* CK1 paralogs (Yck2, Yck22, and Hrr25), with Hog1 as the only off-target protein kinase across the entire fungal proteome.

**Figure 3.**
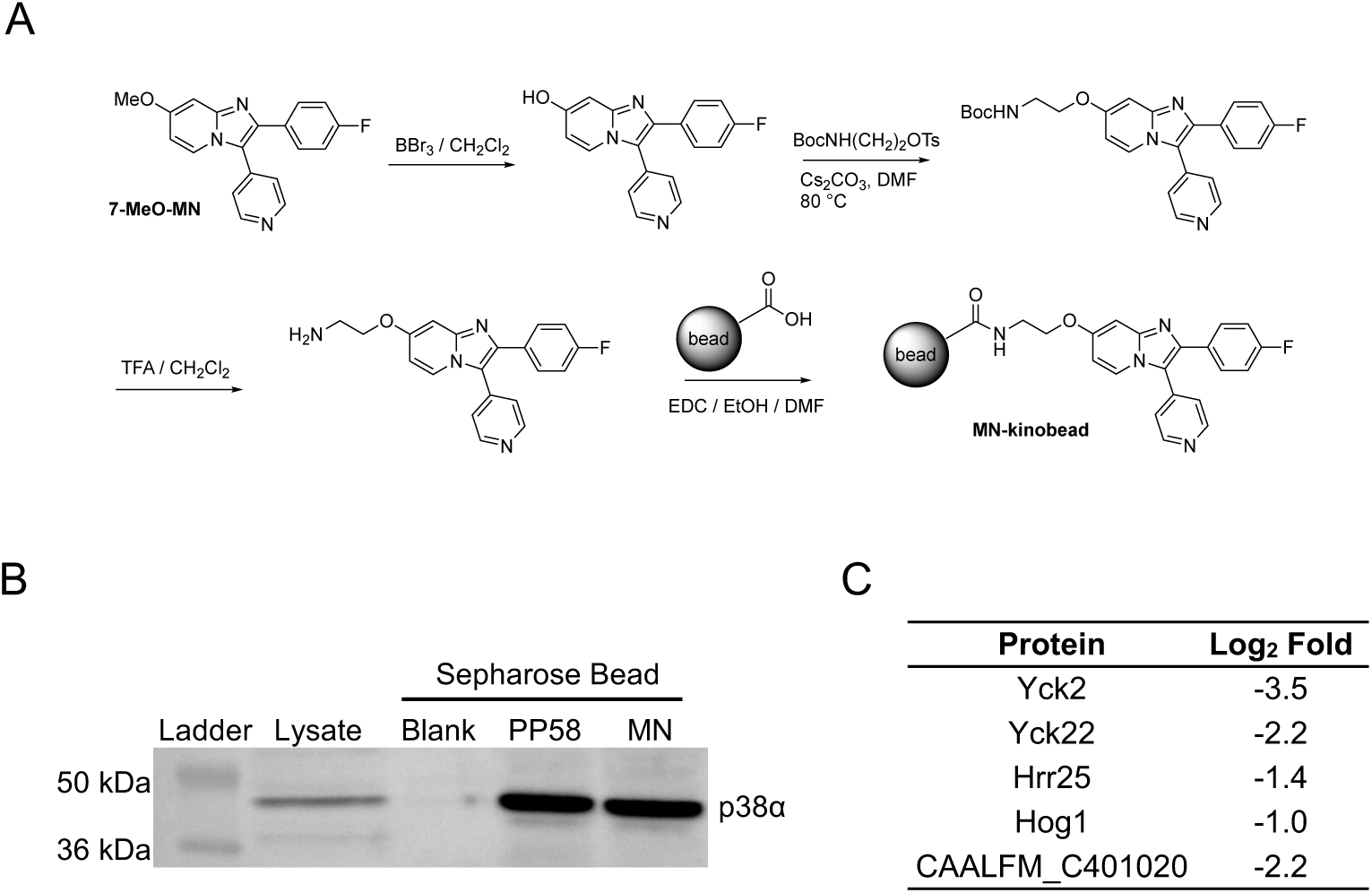
MN-kinobead pulldown experiment. **A.** Synthetic scheme for the MN-kinobead. **B.** Western blot for p38α following pull down from HEK293 cell lysate. **C.** *C. albicans* proteins competed by MN (10 μM) following MN-kinobead pull down from *C. albicans* cell lysate.

To further characterize the kinome-wide selectivity of YK and MN, additional profiling was performed using human cell lysates. MIB/MS competition assays employing HEK293 cell lysates demonstrated that YK and MN engaged eight and seven human protein kinases, respectively, out of a total of 269 identified by MIB/MS. Six of the human kinases were engaged by both compounds (CK1α, CK1δ, CK1ε, p38α, PKN3, RIPK2) (Figure 4A and B, File S6). CK1γ2 and ALK4 were engaged by only YK and p38β by only MN. As was observed in the *Candida* lysate, the most robustly competed kinases in human cell lysate were three CK1 paralogs and p38α. To demonstrate in cell target engagement, NanoBRET assays were performed with YK and MN against a panel of 192 human protein kinases. The NanoBRET assay utilizes a nano-luciferase (Nluc)-tagged kinase in live cells, allowing for robust, quantitative kinase inhibitor profiling in cells for a variety of kinase targets^50^. Notably, 88 of the kinases screened by NanoBRET were not captured from HEK293 cell lysate by MIB (Figure 4C), and thus, provide additional depth to the kinome-wide profiling. In the NanoBRET target engagement assays (Figure 4D), YK (1 µM) demonstrated >70% occupancy of p38α and CK1α while MN (1 µM) showed >70% occupancy of only p38α. YK also showed 40–70% occupancy of NLK, RIPK2, and GAK, with MN showing engagement of CK1α and NLK at these intermediate levels. Notably, the human kinases NLK and GAK were not captured from HEK293 cell lysate by the MIB matrix, suggesting that these proteins were not expressed at detectable levels in the cells.

**Figure 4.**
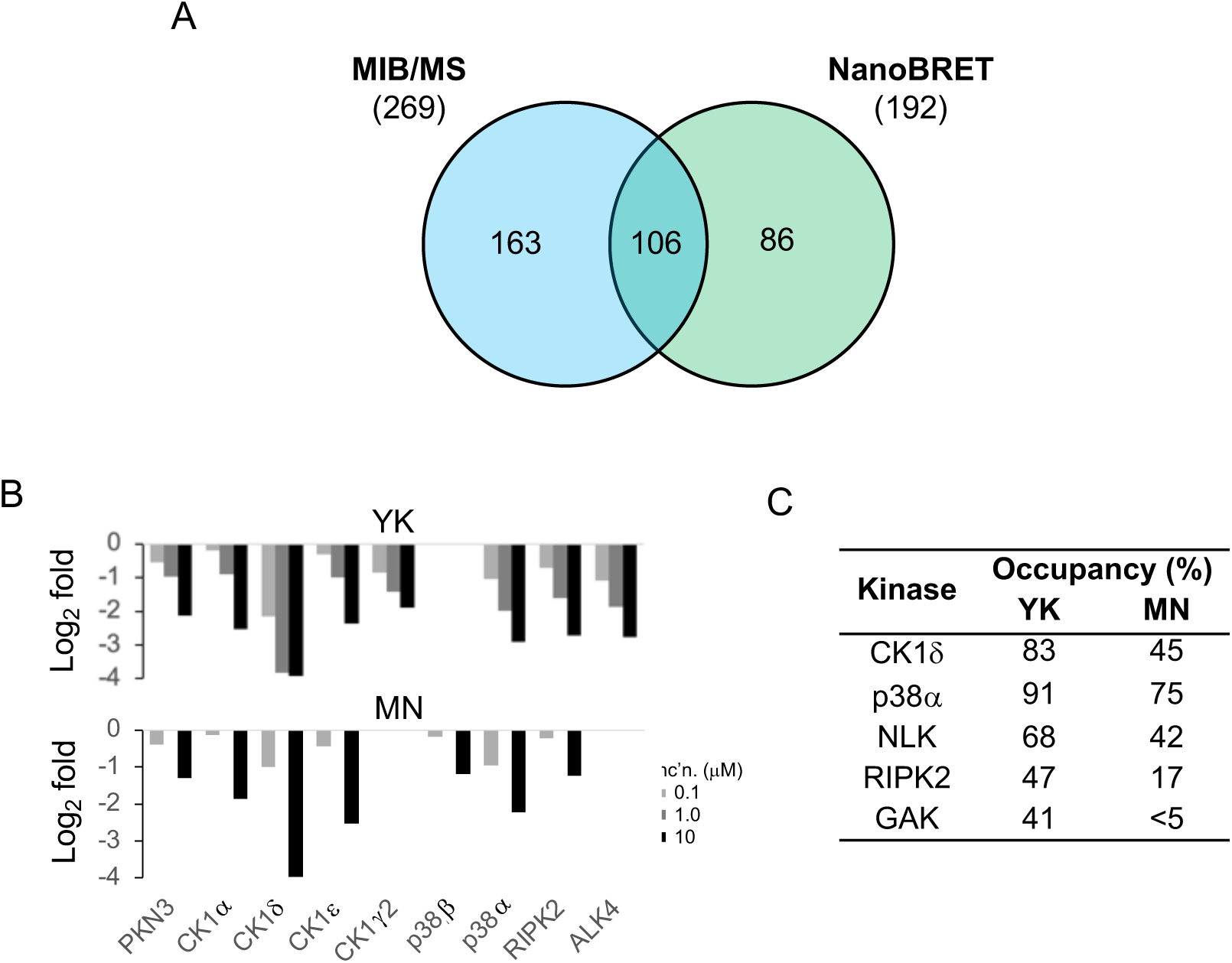
YK and MN selectivity across the human kinome. **A.** Number of human kinases assayed by MIB/MS in blue (269) and NanoBRET (192) in green. A total of 355 human kinases are assayed by the two techniques combined. **B.** Human kinases showing significant dose dependent competition by MIB/MS for either YK or MN. **C.** Target engagement of human kinases expressed as percent occupancy using NanoBRET by 1.0 μM of YK or MN.

In summary, YK and MN showed exceptionally high kinome-wide selectivity by MIB/MS profiling, with engagement of only four homologous kinases across the fungal and human proteome. This high selectivity was confirmed by NanoBRET using in cell target engagement assays. The only non-CK1 family kinase targeted by these inhibitors was MAPK14 (Hog1/p38α). The identification of Hog1 as an off-target kinase was not entirely surprising as GW was designed as a selective human p38α inhibitor^45^. However, it was remarkable that Hog1 was the sole off-target kinase of YK and MN across the entire *Candida* kinome. Given the therapeutic potential of Yck2 inhibitors as single agent antifungals, we sought experimental validation of whether it was possible to identify Yck2 family inhibitors with additional selectivity over Hog1. Our approach leveraged the high homology that was observed between the eukaryotic kinomes in our MIB/MS profiling experiments by mining the vast knowledgebase of human kinase inhibitors for CK1 inhibitors that were reported to be selective over p38α. Sixteen publications detailing kinome-wide screens of ∼2500 unique inhibitors, corresponding to >1 million inhibitor-kinase interactions were triaged (Figure 5A, File S7). Hits were filtered for engagement of any CK1 paralog (IC_50_ < 1 µM or > 90% inhibition at 1 µM) to yield 179 annotations and 150 unique compounds. Seventy-one of these inhibitors originated from our open source PKIS and PKIS2 collections (Figure 5B)^51,52^. An additional 15 CK1 inhibitors were identified by the Küster laboratory in their independent kinobead experiments^39,53^. CK1 inhibitors were also selected from online databases maintained by the Chemical Probes Portal, Millipore, and MedChemExpress^54–56^. 37 of the compounds identified contained aminopyrimidine hinge-binding motifs (Figure 5C). A large portion of these CK1 inhibitors were also phenoxypyrimidines (19) and pyridines (23), but many others had high structural diversity. After filtering for reported inactivity against p38α, thirteen compounds were selected for screening on the fungal kinases as potential Yck2 inhibitors with selectivity over Hog1.

**Figure 5.**
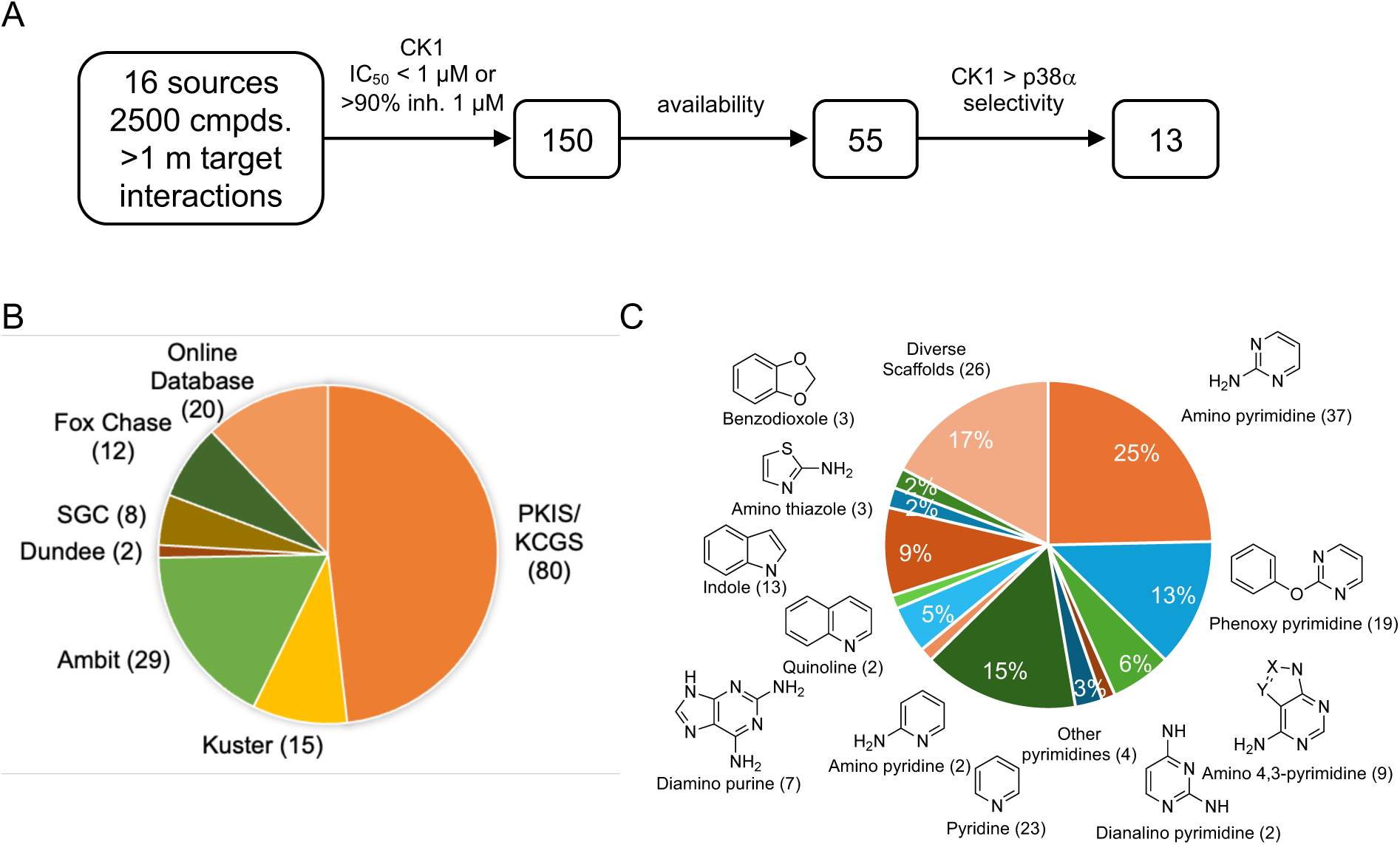
Repurposing of CK1 as potential Yck2 inhibitors. **A.** Flowchart of compound triage for candidate Yck2 inhibitors. **B.** Source of CK1 inhibitor screening data. **C.** CK1 inhibitor chemotypes

GW, YK, MN, and the thirteen candidate inhibitors were assayed for inhibition of *C*. *albicans* Yck2 kinase activity and counter-screened against human CK1α and p38α using biochemical assays employing the purified enzymes (Table 1). In-cell target engagement for CK1 paralogs was confirmed using established NanoBRET assays^50^. To enable a Yck2 NanoBRET assay, a custom Bodipy tracer was synthesized from MN using the chemistry developed for the MN-kinobead (Figure 6A). Titration of MN against NLuc-Yck2 using the MN-Bodipy tracer established 500 nM as the optimal concentration for the NanoBRET assay (Figure 6B).

**Figure 6.**
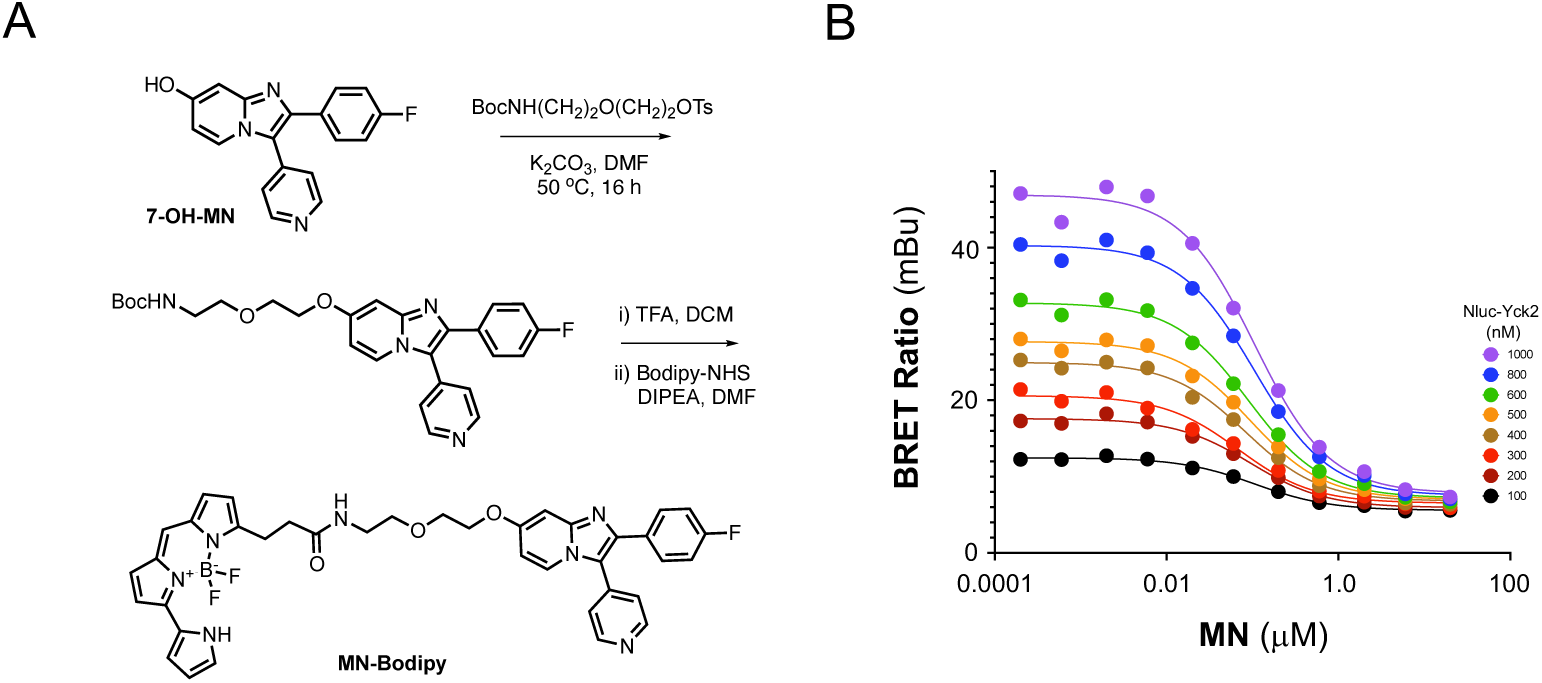
Yck2 NanoBRET assay. A. Synthesis of the MN-Bodipy tracer. B. Tracer titration of MN-Bodipy on NLuc-Yck2.

**Table 1.**
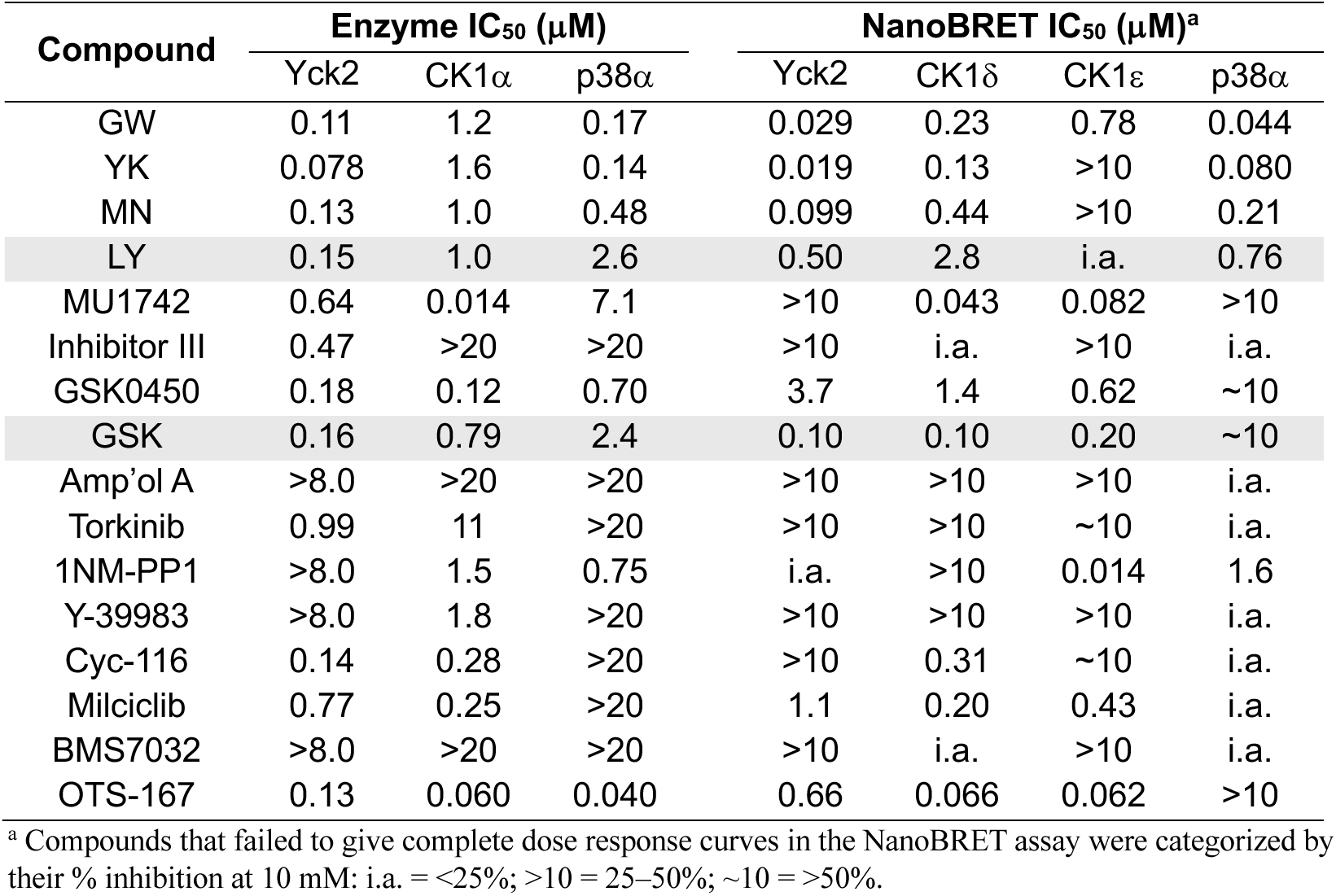
Identification of new Yck2 chemotypes.

GW, YK, and MN showed sub-micromolar inhibition of Yck2 with 8- to 20-fold selectivity over CK1α but less than 3-fold selectivity over p38α. In contrast, many of the thirteen candidate inhibitors showed high selectivity for Yck2 over p38α. Nine inhibitors (LY364947, MU1742, CDK1/2 Inhibitor III, GSK150450B, GSK507358A, Torkinib, Cyc-116, Milciclib, OTS-167) demonstrated IC_50_ < 1 µM for inhibition of the Yck2 enzyme, with three of these (LY364947, GSK507358A, OTS-167) showed sub-micromolar engagement of Yck2 in cells using the newly enabled NanoBRET assay. Ten of the inhibitors were inactive against the p38α enzyme at 1 µM with the remaining three having reduced activity compared to YK and MN. LY364947 (LY) and GSK507358A (GSK) were chosen for further study based on their selectivity across the human kinome and potent Yck2 target engagement, respectively. LY was first reported as a selective ALK5 inhibitor^52,57,58^, while GSK was described an AKT inhibitor^59^.

To determine the fungal kinome profile of LY and GSK, MIB/MS competition assays were conducted using *C. albicans* cell lysate across a dose range of inhibitor from 0.1–10 µM. Sixty-five fungal protein kinases were captured in the MIB/MS competition assay (File S8). LY showed remarkably selective target engagement of only Yck2 and its paralogs, Yck22 and Hrr25 no activity on Hog1 (Figure 7A). While GSK also displayed dose-dependent engagement of Yck2 it had cross-reactivity with Hog1 and 30 additional *Candida* protein kinases at 10 µM (Figure 7B). Thus, the MIB/MS profiling demonstrated that LY had the superior kinase selectivity profile with engagement of only Yck2 and its two paralogs across entire fungal protein kinome.

**Figure 7.**
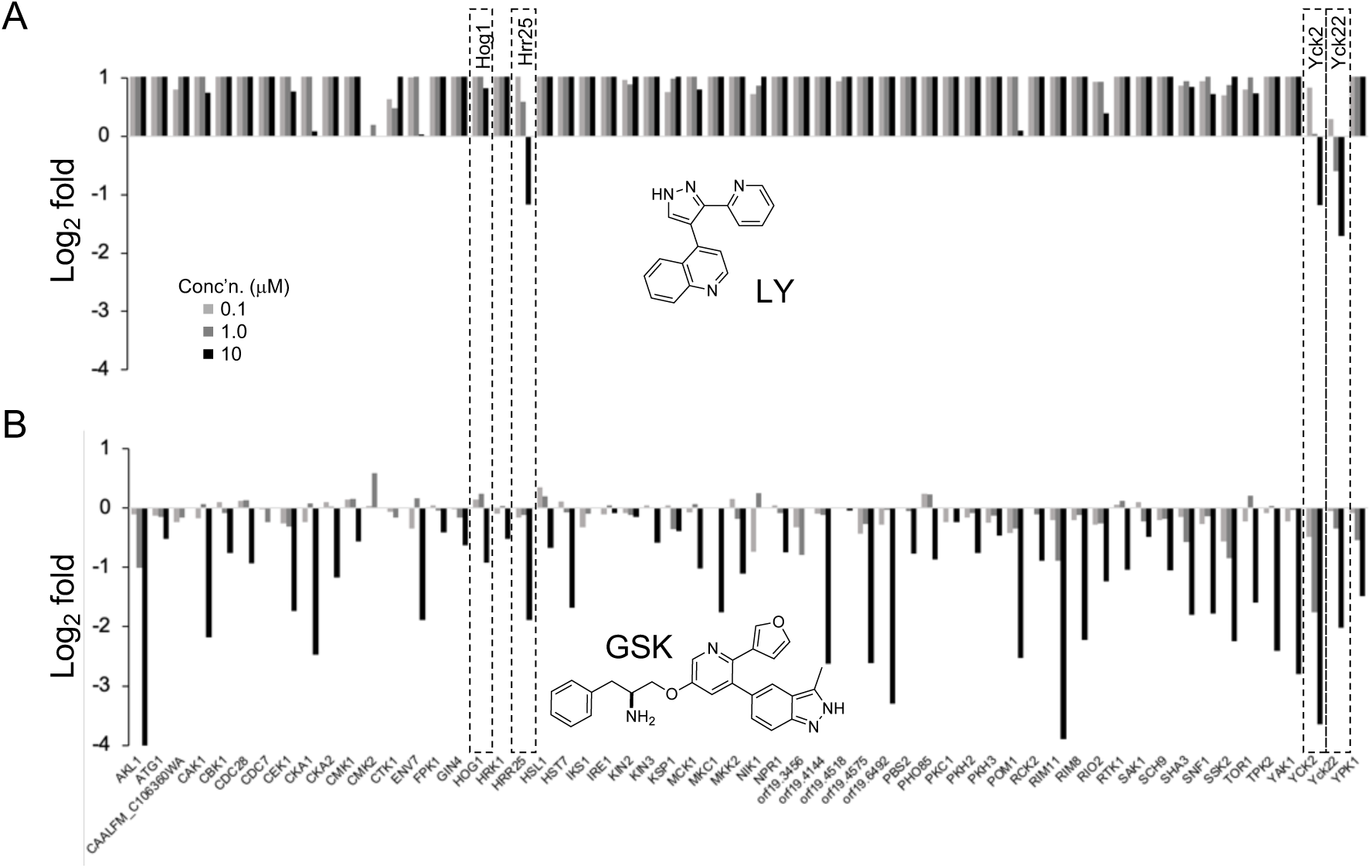
MIB/MS competition assay with LY and GSK. **A.** MIB/MS profile of LY. **B.** MIB/MS profile of GSK.

To determine if selective inhibition of the Yck2 family (without Hog1) would translate to whole cell antifungal activity, LY was tested in two-fold dose-response assays against the wild-type CaSS1 strain of *C. albicans* in RPMI medium at 37 °C with 5% CO_2_ continuous infusion, conditions under which Yck2 is required for growth ^21^. However, under these growth conditions, LY was only weakly active at the top 100 µM concentration (Figure 8A). To confirm that this antifungal activity was dependent on Yck2, LY was retested against a mutant *C. albicans* strain containing only a single allele of *YCK2* that had been placed under the control of a doxycycline (DOX)-repressible promoter (*tetO-YCK2/yck2Δ*). With this engineered strain, in the absence of DOX, robust inhibition of fungal growth was obtained at the 100 μM dose of LY (Figure 8B). Addition of DOX to further reduce the gene dosage of *YCK2* resulted in hypersensitivity of *C. albicans* growth to LY, consistent with inhibition of the Yck2 as the mechanism of antifungal activity. To probe whether the relatively poor antifungal activity of LY against wild-type cells might be due to low intracellular levels of the inhibitor, additional experiments were performed using two other mutant *C. albicans* stains. Using a strain in which four fungal efflux pumps had been deleted (*cdr1Δ/Δ, cdr2Δ/Δ, mdr1Δ/Δ, flu1Δ/Δ*), no increase was observed in the activity of LY (Figure 8C). However, in a *C. albicans* stain where cell membrane integrity was disrupted by deletion of the ERG6 sterol methyltransferase gene upon addition of DOX (*tetO-ERG6/erg6Δ*), LY demonstrated potent antifungal activity (Figure 8D). Therefore, it appeared that whole cell antifungal activity of LY was limited primarily by its poor cell penetration. In summary, LY was identified by MIB profiling as a promising chemotype of Yck2 family selective fungal kinase inhibitors. The whole cell antifungal activity of LY was dependent on YCK2 gene dosage, demonstrating its on target activity. However, the antifungal efficacy of LY was limited by poor whole cell penetration that will require further optimization to enhance its therapeutic utility.

**Figure 8.**
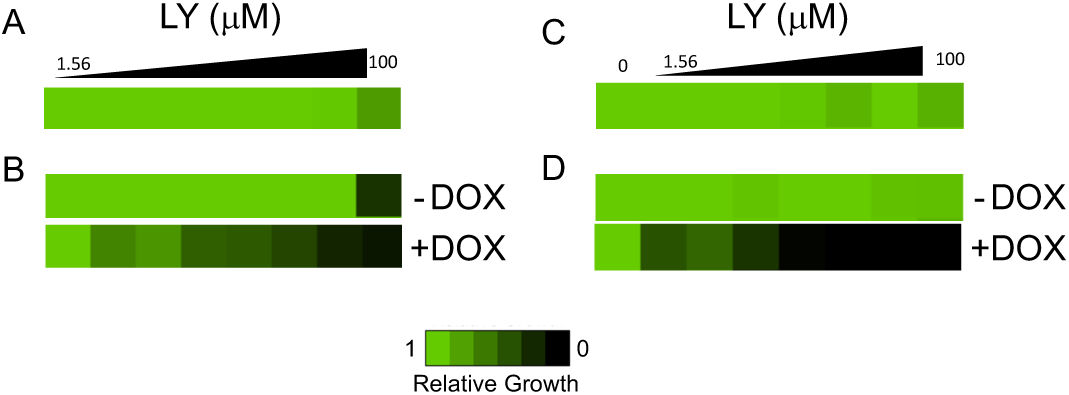
*C. Albicans* antifungal activity of LY. **A.** Wild-type CaSS1. **B**. *tetO-YCK2/yck2τι*. **C**. *cdr1τι/τι, cdr2τι/τι, mdr1τι/τι, flu1τι/τι*. **D**. *tetO-ERG6/erg6τι*. DOX = doxycyclin (5 or 10 μg/mL).

The growing threat of IFI’s and lack of new treatment options has increased the urgency to develop novel, effective, non-toxic antifungals. Many fungal kinases have been identified as essential proteins for cell growth or viability but have yet to be targeted by small molecules inhibitors^15,16^. A handful of fungal kinase inhibitors have been approved only for use as pesticides (e.g. cercosporamide, fludioxonil, and iprodine). In these cases, their mechanism of action was not attributed to kinase inhibition until years, even decades after regulatory approval^16,60–62^. Furthermore, there have been no assays available for measuring inhibitor binding and selectivity across the fungal kinome. We now demonstrate that a MIB matrix built from six promiscuous kinase inhibitors captured 63 of 89 high confidence predicted protein kinases in the fungal kinome, including a majority of kinases across all families. The MIB matrix allows not only rapid characterization of antifungal kinase inhibitors, but also development of chemical probes for exploration of the “dark kinome” of *C. albicans*. According to UniProtKB, less than 30% of the predicted kinases in *C. albicans* have been confirmed experimentally^35^. Our global proteomic analysis is the first experimental validation of the expressed *C. albicans* kinome, totaling 71 protein kinases. Although little information is known about the cellular function of many of these fungal kinases, they may represent a rich source of new chemically tractable antifungal drug targets.

MIB/MS competition assays of the antifungal kinase inhibitors YK and MN showed that they possess remarkable selectivity with engagement of only the Yck2 family and Hog1 across the fungal kinome. These results were confirmed for MN using the bead-immobilized inhibitor, which resulted in the capture of only three Yck2 paralogs and Hog1, plus a single non-kinase protein from *C. albicans* lysate. These kinome profiling results also highlighted the close phylogenetic relationship between the eukaryotic *C. albicans* and *Homo sapiens* kinases likely resulting from the evolution of a common primordial enzyme that utilized ATP as a cofactor^63,64^. Specifically, *C. albicans* Yck2 and the human CK1 paralogs have on average 50% overall sequence identity, a common architecture in the N- and C-lobes, and share many of the key residues utilized in binding ATP and small molecules^65,66^. Using this insight, we leveraged the extensive landscape of human kinase inhibitor knowledge to repurpose CK1 inhibitors with selectivity over p38α as candidate Yck2 inhibitors with selectivity over Hog1. LY emerged as the most promising chemotype that met the criteria of fungal Yck2 kinase inhibition with Hog1 selectivity. Although LY demonstrated on target antifungal activity, it requires additional optimization to improve fungal cell penetration. These future studies will be aided by the availability of MIB/MS profiling to measure fungal kinome selectivity for new drug candidates.

## ACKNOWLEDGEMENTS

The Structural Genomics Consortium (SGC) is a registered charity (no: 1097737) that receives funds from Bayer AG, Boehringer Ingelheim, Bristol Myers Squibb, Genentech, Genome Canada, through Ontario Genomics Institute [OGI-196], EU/EFPIA/OICR/McGill/KTH/Diamond Innovative Medicines Initiative 2 Joint Under-taking [EUbOPEN grant 875510], Janssen, Merck KGaA (also known as EMD in Canada and the US), Pfizer, and Takeda. The research reported in this publication was supported by NIH grants R01AI162789-01 and R01GM138520-04. Research conducted by the UNC Metabolomics and Proteomics Core Facility was supported in part by NIH grant P30CA016086-34. Additional infrastructure support was provided in part by the NC Biotech Center Institutional Support Grant 2018-IDG-1030 and NIH grant S10OD032476 for upgrading the 500 MHz NMR spectrometer in the UNC Eshelman School of Pharmacy NMR Facility. John Forsberg and Nathaniel I. Nicely of the UNC Protein Expression and Purification Core contributed HEK293 cell pellets for the MIB/MS experiments. We thank Merck & Co and Genome Canada for making the original Candida albicans GRACE mutant collections available.

## REFERENCES

(1) Fisher, M. C.; Gurr, S. J.; Cuomo, C. A.; Blehert, D. S.; Jin, H.; Stukenbrock, E. H.; Stajich, J. E.; Kahmann, R.; Boone, C.; Denning, D. W.; Gow, N. A. R.; Klein, B. S.; Kronstad, J. W.; Sheppard, D. C.; Taylor, J. W.; Wright, G. D.; Heitman, J.; Casadevall, A.; Cowen, L. E. Threats Posed by the Fungal Kingdom to Humans, Wildlife, and Agriculture. mBio 2020, 11 (3). 10.1128/mBio.00449-20.

(2) Denning, D. W. Global Incidence and Mortality of Severe Fungal Disease. Lancet Infect Dis 2024, 24 (7), e428–e438. 10.1016/S1473-3099(23)00692-8.

(3) Brown, G. D.; Denning, D. W.; Levitz, S. M. Tackling Human Fungal Infections. Science (1979) 2012, 336 (6082), 647–647. 10.1126/science.1222236.

(4) W.H.O. WHO Fungal Priority Pathogens List to Guide Research, Development and Public Health Action; 2022. https://www.who.int/publications/i/item/9789240060241 (accessed 2024-07-23).

(5) Papon, N.; Nevez, G.; Le Gal, S.; Vigneau, C.; Robert-Gangneux, F.; Bouchara, J.-P.; Cornely, O. A.; Denning, D. W.; Gangneux, J.-P. Fungal Infections in Transplant Recipients: Pros and Cons of Immunosuppressive and Antimicrobial Treatment. Lancet Microbe 2021, 2 (1), e6–e8. 10.1016/S2666-5247(20)30199-3.

(6) C.D.C. Antimicrobial Resistance Threats in the United States, 2021-2022; 2024. https://www.cdc.gov/antimicrobial-resistance/data-research/threats/update-2022.html (accessed 2024-07-24).

(7) Brown, G. D.; Denning, D. W.; Gow, N. A. R.; Levitz, S. M.; Netea, M. G.; White, T. C. Hidden Killers: Human Fungal Infections. Sci Transl Med 2012, 4 (165). 10.1126/scitranslmed.3004404.

(8) Pfaller, M. A.; Diekema, D. J.; Turnidge, J. D.; Castanheira, M.; Jones, R. N. Twenty Years of the SENTRY Antifungal Surveillance Program: Results for Candida Species From 1997–2016. Open Forum Infect Dis 2019, 6 (Supplement_1), S79–S94. 10.1093/ofid/ofy358.

(9) Pfaller, M. A.; Diekema, D. J. Epidemiology of Invasive Mycoses in North America. Crit Rev Microbiol 2010, 36 (1), 1–53. 10.3109/10408410903241444.

(10) Perlin, D. S.; Rautemaa-Richardson, R.; Alastruey-Izquierdo, A. The Global Problem of Antifungal Resistance: Prevalence, Mechanisms, and Management. Lancet Infect Dis 2017, 17 (12), e383–e392. 10.1016/S1473-3099(17)30316-X.

(11) Moran, M.; Chapman, N.; Abela-Oversteegen, L.; Chowdhary, V.; Doubell, A.; Whittall, C.; Howard, R.; Farrell, P.; Halliday, D.; Hirst, C. NEGLECTED DISEASE RESEARCH AND DEVELOPMENT: THE EBOLA EFFECT; 2015. https://www.policycures.org/wp-content/uploads/2022/11/Y8-GFINDER-full-report-web.pdf (accessed 2024-07-23).

(12) Attwood, M. M.; Fabbro, D.; Sokolov, A. V.; Knapp, S.; Schiöth, H. B. Trends in Kinase Drug Discovery: Targets, Indications and Inhibitor Design. Nat Rev Drug Discov 2021, 20 (11), 839–861. 10.1038/s41573-021-00252-y.

(13) Cohen, P.; Cross, D.; Jänne, P. A. Kinase Drug Discovery 20 Years after Imatinib: Progress and Future Directions. Nat Rev Drug Discov 2021, 20 (7), 551–569. 10.1038/s41573-021-00195-4.

(14) Ardito, F.; Giuliani, M.; Perrone, D.; Troiano, G.; Muzio, L. Lo. The Crucial Role of Protein Phosphorylation in Cell Signaling and Its Use as Targeted Therapy (Review). Int J Mol Med 2017, 40 (2), 271–280. 10.3892/ijmm.2017.3036.

(15) Lee, K.-T.; So, Y.-S.; Yang, D.-H.; Jung, K.-W.; Choi, J.; Lee, D.-G.; Kwon, H.; Jang, J.; Wang, L. L.; Cha, S.; Meyers, G. L.; Jeong, E.; Jin, J.-H.; Lee, Y.; Hong, J.; Bang, S.; Ji, J.-H.; Park, G.; Byun, H.-J.; Park, S. W.; Park, Y.-M.; Adedoyin, G.; Kim, T.; Averette, A. F.; Choi, J.-S.; Heitman, J.; Cheong, E.; Lee, Y.-H.; Bahn, Y.-S. Systematic Functional Analysis of Kinases in the Fungal Pathogen Cryptococcus Neoformans. Nat Commun 2016, 7 (1), 12766. 10.1038/ncomms12766.

(16) Hameed, S.; Rehman, S.; Fatima, Z.; Hameed, S. Protein Kinases as Potential Anticandidal Drug Targets. Frontiers in Bioscience 2020, 25 (8), 4862. 10.2741/4862.

(17) Caplan, T.; Lorente-Macías, Á.; Stogios, P. J.; Evdokimova, E.; Hyde, S.; Wellington, M. A.; Liston, S.; Iyer, K. R.; Puumala, E.; Shekhar-Guturja, T.; Robbins, N.; Savchenko, A.; Krysan, D. J.; Whitesell, L.; Zuercher, W. J.; Cowen, L. E. Overcoming Fungal Echinocandin Resistance through Inhibition of the Non-Essential Stress Kinase Yck2. Cell Chem Biol 2020, 27 (3), 269–282.e5. 10.1016/j.chembiol.2019.12.008.

(18) Elkins, J. M.; Fedele, V.; Szklarz, M.; Abdul Azeez, K. R.; Salah, E.; Mikolajczyk, J.; Romanov, S.; Sepetov, N.; Huang, X.-P.; Roth, B. L.; Al Haj Zen, A.; Fourches, D.; Muratov, E.; Tropsha, A.; Morris, J.; Teicher, B. A.; Kunkel, M.; Polley, E.; Lackey, K. E.; Atkinson, F. L.; Overington, J. P.; Bamborough, P.; Müller, S.; Price, D. J.; Willson, T. M.; Drewry, D. H.; Knapp, S.; Zuercher, W. J. Comprehensive Characterization of the Published Kinase Inhibitor Set. Nat Biotechnol 2016, 34 (1), 95–103. 10.1038/nbt.3374.

(19) Wells, C. I.; Al-Ali, H.; Andrews, D. M.; Asquith, C. R. M.; Axtman, A. D.; Dikic, I.; Ebner, D.; Ettmayer, P.; Fischer, C.; Frederiksen, M.; Futrell, R. E.; Gray, N. S.; Hatch, S. B.; Knapp, S.; Lücking, U.; Michaelides, M.; Mills, C. E.; Müller, S.; Owen, D.; Picado, A.; Saikatendu, K. S.; Schröder, M.; Stolz, A.; Tellechea, M.; Turunen, B. J.; Vilar, S.; Wang, J.; Zuercher, W. J.; Willson, T. M.; Drewry, D. H. The Kinase Chemogenomic Set (KCGS): An Open Science Resource for Kinase Vulnerability Identification. Int J Mol Sci 2021, 22 (2), 566. 10.3390/ijms22020566.

(20) Jung, S.-I.; Rodriguez, N.; Irrizary, J.; Liboro, K.; Bogarin, T.; Macias, M.; Eivers, E.; Porter, E.; Filler, S. G.; Park, H. Yeast Casein Kinase 2 Governs Morphology, Biofilm Formation, Cell Wall Integrity, and Host Cell Damage of Candida Albicans. PLoS One 2017, 12 (11), e0187721. 10.1371/journal.pone.0187721.

(21) Cowen, L.; Puumala, E; Nandakumar, M; Yiu, B; Stogios, P; Strickland, B; Zarnowski, R; Wang, X; Williams, N; Savchenko, A; Andes, D; Robbins, N; Whitesell, L; Willson, T. Structure-guided Optimization of Small Molecules Targeting the Yeast Casein Kinase, Yck2, as a Therapeutic Strategy to Combat Candida albicans. Res Sq [Preprint] 2025, Jan 08, rs.3.rs-5524306, 10.21203/rs.3.rs-5524306/v1.

(22) Chan, W. C.; Sharifzadeh, S.; Buhrlage, S. J.; Marto, J. A. Chemoproteomic Methods for Covalent Drug Discovery. Chem Soc Rev 2021, 50 (15), 8361–8381. 10.1039/D1CS00231G.

(23) Gao, Y.; Ma, M.; Li, W.; Lei, X. Chemoproteomics, A Broad Avenue to Target Deconvolution. Advanced Science 2024, 11 (8). 10.1002/advs.202305608.

(24) Radu, M.; Chernoff, J. Recent Advances in Methods to Assess the Activity of the Kinome. F1000Res 2017, 6, 1004. 10.12688/f1000research.10962.1.

(25) Reinecke, M.; Heinzlmeir, S.; Wilhelm, M.; Médard, G.; Klaeger, S.; Kuster, B. Kinobeads: A Chemical Proteomic Approach for Kinase Inhibitor Selectivity Profiling and Target Discovery; Plowright, A. T., Ed.; 2019; pp 97–130. 10.1002/9783527818242.ch4.

(26) Duncan, J. S.; Whittle, M. C.; Nakamura, K.; Abell, A. N.; Midland, A. A.; Zawistowski, J. S.; Johnson, N. L.; Granger, D. A.; Jordan, N. V.; Darr, D. B.; Usary, J.; Kuan, P.-F.; Smalley, D. M.; Major, B.; He, X.; Hoadley, K. A.; Zhou, B.; Sharpless, N. E.; Perou, C. M.; Kim, W. Y.; Gomez, S. M.; Chen, X.; Jin, J.; Frye, S. V.; Earp, H. S.; Graves, L. M.; Johnson, G. L. Dynamic Reprogramming of the Kinome in Response to Targeted MEK Inhibition in Triple-Negative Breast Cancer. Cell 2012, 149 (2), 307–321. 10.1016/j.cell.2012.02.053.

(27) Cousins, E. M.; Goldfarb, D.; Yan, F.; Roques, J.; Darr, D.; Johnson, G. L.; Major, M. B. Competitive Kinase Enrichment Proteomics Reveals That Abemaciclib Inhibits GSK3β and Activates WNT Signaling. Molecular Cancer Research 2018, 16 (2), 333–344. 10.1158/1541-7786.MCR-17-0468.

(28) Graves, L. M.; Duncan, J. S.; Whittle, M. C.; Johnson, G. L. The Dynamic Nature of the Kinome. Biochemical Journal 2013, 450 (1), 1–8. 10.1042/BJ20121456.

(29) Zawistowski, J. S.; Graves, L. M.; Johnson, G. L. Assessing Adaptation of the Cancer Kinome in Response to Targeted Therapies. Biochem Soc Trans 2014, 42 (4), 765–769. 10.1042/BST20130274.

(30) Fritch, E. J.; Mordant, A. L.; Gilbert, T. S. K.; Wells, C. I.; Yang, X.; Barker, N. K.; Madden, E. A.; Dinnon, K. H.; Hou, Y. J.; Tse, L. V.; Castillo, I. N.; Sims, A. C.; Moorman, N. J.; Lakshmanane, P.; Willson, T. M.; Herring, L. E.; Graves, L. M.; Baric, R. S. Investigation of the Host Kinome Response to Coronavirus Infection Reveals PI3K/MTOR Inhibitors as Betacoronavirus Antivirals. J Proteome Res 2023, 22 (10), 3159–3177. 10.1021/acs.jproteome.3c00182.

(31) Krulikas, L. J.; McDonald, I. M.; Lee, B.; Okumu, D. O.; East, M. P.; Gilbert, T. S. K.; Herring, L. E.; Golitz, B. T.; Wells, C. I.; Axtman, A. D.; Zuercher, W. J.; Willson, T. M.; Kireev, D.; Yeh, J. J.; Johnson, G. L.; Baines, A. T.; Graves, L. M. Application of Integrated Drug Screening/Kinome Analysis to Identify Inhibitors of Gemcitabine-Resistant Pancreatic Cancer Cell Growth. SLAS Discovery 2018, 23 (8), 850–861. 10.1177/2472555218773045.

(32) Stewart, H.; Grinfeld, D.; Hagedorn, B.; Ostermann, R.; Makarov, A.; Hock, C. Proof of Principle for Enhanced Resolution Multi-Pass Methods for the Astral Analyzer. Int J Mass Spectrom 2024, 498, 117203. 10.1016/j.ijms.2024.117203.

(33) Heil, L. R.; Damoc, E.; Arrey, T. N.; Pashkova, A.; Denisov, E.; Petzoldt, J.; Peterson, A. C.; Hsu, C.; Searle, B. C.; Shulman, N.; Riffle, M.; Connolly, B.; MacLean, B. X.; Remes, P. M.; Senko, M. W.; Stewart, H. I.; Hock, C.; Makarov, A. A.; Hermanson, D.; Zabrouskov, V.; Wu, C. C.; MacCoss, M. J. Evaluating the Performance of the Astral Mass Analyzer for Quantitative Proteomics Using Data-Independent Acquisition. J Proteome Res 2023, 22 (10), 3290–3300. 10.1021/acs.jproteome.3c00357.

(34) Guzman, U. H.; Martinez-Val, A.; Ye, Z.; Damoc, E.; Arrey, T. N.; Pashkova, A.; Renuse, S.; Denisov, E.; Petzoldt, J.; Peterson, A. C.; Harking, F.; Østergaard, O.; Rydbirk, R.; Aznar, S.; Stewart, H.; Xuan, Y.; Hermanson, D.; Horning, S.; Hock, C.; Makarov, A.; Zabrouskov, V.; Olsen, J. V. Ultra-Fast Label-Free Quantification and Comprehensive Proteome Coverage with Narrow-Window Data-Independent Acquisition. Nat Biotechnol 2024. 10.1038/s41587-023-02099-7.

(35) Proteomes - Candida albicans (strain SC5314/ATCC MYA-2876) (Yeast). UniProt. https://www.uniprot.org/proteomes/UP000000559 (accessed 2024-07-24).

(36) Tyanova, S.; Temu, T.; Sinitcyn, P.; Carlson, A.; Hein, M. Y.; Geiger, T.; Mann, M.; Cox, J. The Perseus Computational Platform for Comprehensive Analysis of (Prote)Omics Data. Nat Methods 2016, 13 (9), 731–740. 10.1038/nmeth.3901.

(37) Candida Genome Database. http://www.candidagenome.org/news_archive.shtml (accessed 2024-07-24).

(38) Das, S.; Bhuyan, R.; Goswami, A. M.; Saha, T. Kinome Analyses of Candida Albicans, C. Parapsilosis and C. Tropicalis Enable Novel Kinases as Therapeutic Drug Targets in Candidiasis. Gene 2021, 780, 145530. 10.1016/j.gene.2021.145530.

(39) Klaeger, S.; Heinzlmeir, S.; Wilhelm, M.; Polzer, H.; Vick, B.; Koenig, P.-A.; Reinecke, M.; Ruprecht, B.; Petzoldt, S.; Meng, C.; Zecha, J.; Reiter, K.; Qiao, H.; Helm, D.; Koch, H.; Schoof, M.; Canevari, G.; Casale, E.; Depaolini, S. R.; Feuchtinger, A.; Wu, Z.; Schmidt, T.; Rueckert, L.; Becker, W.; Huenges, J.; Garz, A.-K.; Gohlke, B.-O.; Zolg, D. P.; Kayser, G.; Vooder, T.; Preissner, R.; Hahne, H.; Tõnisson, N.; Kramer, K.; Götze, K.; Bassermann, F.; Schlegl, J.; Ehrlich, H.-C.; Aiche, S.; Walch, A.; Greif, P. A.; Schneider, S.; Felder, E. R.; Ruland, J.; Médard, G.; Jeremias, I.; Spiekermann, K.; Kuster, B. The Target Landscape of Clinical Kinase Drugs. Science (1979) 2017, 358 (6367). 10.1126/science.aan4368.

(40) Klaeger, S.; Gohlke, B.; Perrin, J.; Gupta, V.; Heinzlmeir, S.; Helm, D.; Qiao, H.; Bergamini, G.; Handa, H.; Savitski, M. M.; Bantscheff, M.; Médard, G.; Preissner, R.; Kuster, B. Chemical Proteomics Reveals Ferrochelatase as a Common Off-Target of Kinase Inhibitors. ACS Chem Biol 2016, 11 (5), 1245–1254. 10.1021/acschembio.5b01063.

(41) Connie Truong. Functional Analysis of Yeast Casein Kinase 22 in Vacuolar Trafficking of Candida Ablicans. M.S., California State University, Los Angeles, 2018.

(42) Broxton, C. N.; He, B.; Bruno, V. M.; Culotta, V. C. A Role for Candida Albicans Superoxide Dismutase Enzymes in Glucose Signaling. Biochem Biophys Res Commun 2018, 495 (1), 814–820. 10.1016/j.bbrc.2017.11.084.

(43) Lee, Y.; Liston, S. D.; Lee, D.; Robbins, N.; Cowen, L. E. Functional Analysis of the Candida Albicans Kinome Reveals Hrr25 as a Regulator of Antifungal Susceptibility. iScience 2022, 25 (6), 104432. 10.1016/j.isci.2022.104432.

(44) Correia, I.; Wilson, D.; Hube, B.; Pla, J. Characterization of a Candida Albicans Mutant Defective in All MAPKs Highlights the Major Role of Hog1 in the MAPK Signaling Network. Journal of Fungi 2020, 6 (4), 230. 10.3390/jof6040230.

(45) Cheung, M.; Harris, P. A.; Badiang, J. G.; Peckham, G. E.; Chamberlain, S. D.; Alberti, M. J.; Jung, D. K.; Harris, S. S.; Bramson, N. H.; Epperly, A. H.; Stimpson, S. A.; Peel, M. R. The Identification of Pyrazolo[1,5-a]Pyridines as Potent P38 Kinase Inhibitors. Bioorg Med Chem Lett 2008, 18 (20), 5428–5430. 10.1016/j.bmcl.2008.09.040.

(46) MacAlpine, J.; Liu, Z.; Hossain, S.; Whitesell, L.; Robbins, N.; Cowen, L. E. DYRK-Family Kinases Regulate *Candida Albicans* Morphogenesis and Virulence through the Ras1/PKA Pathway. mBio 2023, 14 (6). 10.1128/mbio.02183-23.

(47) MacAlpine, J.; Liu, Z.; Hossain, S.; Whitesell, L.; Robbins, N.; Cowen, L. E. DYRK-Family Kinases Regulate *Candida Albicans* Morphogenesis and Virulence through the Ras1/PKA Pathway. mBio 2023, 14 (6). 10.1128/mbio.02183-23.

(48) UniProt. A0A1D8PL84 · A0A1D8PL84_CANAL. https://www.uniprot.org/uniprotkb/A0A1D8PL84/entry.

(49) Candida Genome Database. C. albicans C4_01020C Summary. http://www.candidagenome.org/cgi-bin/locus.pl?locus=C4_01020C_A.

(50) Vasta, J. D.; Corona, C. R.; Wilkinson, J.; Zimprich, C. A.; Hartnett, J. R.; Ingold, M. R.; Zimmerman, K.; Machleidt, T.; Kirkland, T. A.; Huwiler, K. G.; Ohana, R. F.; Slater, M.; Otto, P.; Cong, M.; Wells, C. I.; Berger, B.-T.; Hanke, T.; Glas, C.; Ding, K.; Drewry, D. H.; Huber, K. V. M.; Willson, T. M.; Knapp, S.; Müller, S.; Meisenheimer, P. L.; Fan, F.; Wood, K. V.; Robers, M. B. Quantitative, Wide-Spectrum Kinase Profiling in Live Cells for Assessing the Effect of Cellular ATP on Target Engagement. Cell Chem Biol 2018, 25 (2), 206–214.e11. 10.1016/j.chembiol.2017.10.010.

(51) Drewry, D.; Willson, T.; Zuercher, W. Seeding Collaborations to Advance Kinase Science with the GSK Published Kinase Inhibitor Set (PKIS). Curr Top Med Chem 2014, 14 (3), 340–342. 10.2174/1568026613666131127160819.

(52) Drewry, D. H.; Wells, C. I.; Andrews, D. M.; Angell, R.; Al-Ali, H.; Axtman, A. D.; Capuzzi, S. J.; Elkins, J. M.; Ettmayer, P.; Frederiksen, M.; Gileadi, O.; Gray, N.; Hooper, A.; Knapp, S.; Laufer, S.; Luecking, U.; Michaelides, M.; Müller, S.; Muratov, E.; Denny, R. A.; Saikatendu, K. S.; Treiber, D. K.; Zuercher, W. J.; Willson, T. M. Progress towards a Public Chemogenomic Set for Protein Kinases and a Call for Contributions. PLoS One 2017, 12 (8), e0181585. 10.1371/journal.pone.0181585.

(53) Reinecke, M.; Brear, P.; Vornholz, L.; Berger, B.-T.; Seefried, F.; Wilhelm, S.; Samaras, P.; Gyenis, L.; Litchfield, D. W.; Médard, G.; Müller, S.; Ruland, J.; Hyvönen, M.; Wilhelm, M.; Kuster, B. Chemical Proteomics Reveals the Target Landscape of 1,000 Kinase Inhibitors. Nat Chem Biol 2024, 20 (5), 577–585. 10.1038/s41589-023-01459-3.

(54) Antolin, A. A.; Sanfelice, D.; Crisp, A.; Villasclaras Fernandez, E.; Mica, I. L.; Chen, Y.; Collins, I.; Edwards, A.; Müller, S.; Al-Lazikani, B.; Workman, P. The Chemical Probes Portal: An Expert Review-Based Public Resource to Empower Chemical Probe Assessment, Selection and Use. Nucleic Acids Res 2023, 51 (D1), D1492–D1502. 10.1093/nar/gkac909.

(55) Millipore Sigma Home Page. https://www.sigmaaldrich.com/US/en.

(56) MedChemExpress Home Page. https://www.medchemexpress.com/?srsltid=AfmBOoo3M51MeTDFVLD9oKD2o2WkjITVvrEtou01vY2G4acUIN_M1ezK.

(57) Sawyer, J. S.; Anderson, B. D.; Beight, D. W.; Campbell, R. M.; Jones, M. L.; Herron, D. K.; Lampe, J. W.; McCowan, J. R.; McMillen, W. T.; Mort, N.; Parsons, S.; Smith, E. C. R.; Vieth, M.; Weir, L. C.; Yan, L.; Zhang, F.; Yingling, J. M. Synthesis and Activity of New Aryl- and Heteroaryl-Substituted Pyrazole Inhibitors of the Transforming Growth Factor-β Type I Receptor Kinase Domain. J Med Chem 2003, 46 (19), 3953–3956. 10.1021/jm0205705.

(58) Singh, J.; Chuaqui, C. E.; Boriack-Sjodin, P. A.; Lee, W.-C.; Pontz, T.; Corbley, M. J.; Cheung, H.-K.; Arduini, R. M.; Mead, J. N.; Newman, M. N.; Papadatos, J. L.; Bowes, S.; Josiah, S.; Ling, L. E. Successful Shape-Based Virtual Screening: The Discovery of a Potent Inhibitor of the Type I TGFβ Receptor Kinase (TβRI). Bioorg Med Chem Lett 2003, 13 (24), 4355–4359. 10.1016/j.bmcl.2003.09.028.

(59) Lin, H.; Yamashita, D. S.; Zeng, J.; Xie, R.; Wang, W.; Nidarmarthy, S.; Luengo, J. I.; Rhodes, N.; Knick, V. B.; Choudhry, A. E.; Lai, Z.; Minthorn, E. A.; Strum, S. L.; Wood, E. R.; Elkins, P. A.; Concha, N. O.; Heerding, D. A. 2,3,5-Trisubstituted Pyridines as Selective AKT Inhibitors—Part I: Substitution at 2-Position of the Core Pyridine for ROCK1 Selectivity. Bioorg Med Chem Lett 2010, 20 (2), 673–678. 10.1016/j.bmcl.2009.11.064.

(60) Sussman, A.; Huss, K.; Chio, L.-C.; Heidler, S.; Shaw, M.; Ma, D.; Zhu, G.; Campbell, R. M.; Park, T.-S.; Kulanthaivel, P.; Scott, J. E.; Carpenter, J. W.; Strege, M. A.; Belvo, M. D.; Swartling, J. R.; Fischl, A.; Yeh, W.-K.; Shih, C.; Ye, X. S. Discovery of Cercosporamide, a Known Antifungal Natural Product, as a Selective Pkc1 Kinase Inhibitor through High-Throughput Screening. Eukaryot Cell 2004, 3 (4), 932–943. 10.1128/EC.3.4.932-943.2004.

(61) Ochiai, N.; Fujimura, M.; Oshima, M.; Motoyama, T.; Ichiishi, A.; Yamada-Okabe, H.; Yamaguchi, I. Effects of Iprodione and Fludioxonil on Glycerol Synthesis and Hyphal Development in *Candida Albicans*. Biosci Biotechnol Biochem 2002, 66 (10), 2209–2215. 10.1271/bbb.66.2209.

(62) Singh, A.; Sharma, S.; Khuller, G. K. CAMP Regulates Vegetative Growth and Cell Cycle in Candida Albicans. Mol Cell Biochem 2007, 304 (1–2), 331–341. 10.1007/s11010-007-9516-4.

(63) Bougnoux, M.-E.; Aanensen, D. M.; Morand, S.; Théraud, M.; Spratt, B. G.; d’Enfert, C. Multilocus Sequence Typing of Candida Albicans: Strategies, Data Exchange and Applications. Infection, Genetics and Evolution 2004, 4 (3), 243–252. 10.1016/j.meegid.2004.06.002.

(64) McManus, B. A.; Coleman, D. C. Molecular Epidemiology, Phylogeny and Evolution of Candida Albicans. Infection, Genetics and Evolution 2014, 21, 166–178. 10.1016/j.meegid.2013.11.008.

(65) Stogios, P. J.; Evdokimona, E.; Di Leo, R.; Savchenko, A.; Joachimiak, A.; Satchell, K. J. F. Crystal structure of Yck2 from Candida albicans, apoenzyme. 10.2210/pdb6U69/pdb.

(66) Ursu, A.; Illich, D. J.; Takemoto, Y.; Porfetye, A. T.; Zhang, M.; Brockmeyer, A.; Janning, P.; Watanabe, N.; Osada, H.; Vetter, I. R.; Ziegler, S.; Schoeler, H. R.; Waldmann, H. Human Casein Kinase 1 isoform delta apo (kinase domain). 10.2210/pdb5IH4/pdb.

