## Supporting Information for "Chemoproteomic Profiling of *C. albicans* for Characterization of Anti-fungal Kinase Inhibitors"

**Table of Contents**

| | Page |
|---|---|
| Supplementary materials and methods | S2 |
| Figure S1, Western blot of MN-kinobead | S13 |
| References | S14 |
| Compound characterization | S15 |

**General Chemistry Methods.** All reagents and solvents used were purchased from commercial sources and were used without further purification. NMR spectra were obtained using Bruker 400 MHz spectrometers at room temperature; chemical shifts are expressed in parts per million (ppm, δ units) and are referenced to the residual protons in the deuterated solvent used. Coupling constants are given in units of hertz (Hz). Splitting patterns describe apparent multiplicities and are designated as s (singlet), d (doublet), t (triplet), q (quartet), m (multiplet), and br s (broad singlet), dd (doublet of doublets), ddd (double double doublet), tt (triplet of triplets). The purity of compounds submitted for biological screening was determined to be ≥95% as measured by HPLC. Analytical thin layer chromatography (TLC) was performed on silica gel plates, 200 μm with an F254 indicator. Column chromatography was performed using RediSep Rf preloaded silica gel cartridges on Isolera one Biotage automated purification systems. Samples for high-resolution mass spectrometry were analyzed with a ThermoFisher Q Exactive HF-X (ThermoFisher, Bremen, Germany) mass spectrometer coupled with a Waters Acquity H-class liquid chromatograph system. Samples were introduced by a heated electrospray source (HESI) at a flow rate of 0.3 mL/min. Electrospray source conditions were set as: spray voltage 3.0 kV, sheath gas (nitrogen) 60 arb, auxillary gas (nitrogen) 20 arb, sweep gas (nitrogen) 0 arb, nebulizer temperature 375 °C, capillary temperature 380 °C, RF funnel 45 V. The mass range was set to 150–2000 m/z. All measurements were recorded at a resolution setting of 120,000. Separations were conducted on a Waters Acquity UPLC BEH C18 column (2.1 x 50 mm, 1.7 $\mu$m particle size). LC conditions were set at 95 % water with 0.1% formic acid (A) ramped linearly over 5.0 min to 100% acetonitrile with 0.1% formic acid (B) and held until 6.0 min. At 7.0 min the gradient was switched back to 95% (A) and allowed to re-equilibrate until 9.0 min. Injection volume for all samples was 3 $\mu$L. Analytical LC/MS data was obtained using a Waters Acquity Ultrahigh-performance liquid chromatography (UPLC) system equipped with a photodiode array (PDA) detector using the following method: solvent A = Water + 0.2% FA, solvent B = ACN + 0.1% FA, flow rate = 1 mL/min. The gradient started at 95% A for 0.05 min, ramped up to 100% B over 2 min and held for an additional minute at this concentration, before returning to the initial gradient. Compounds were purified by preparative HPLC using an Agilent 1100 equipped with a Phenomenex column (Phenyl-Hexyl, 75 x 30 mm, 5 μm) using the following method: Solvent A: water + 0.05 % TFA; Solvent B: MeOH; flow rate: 70.0 mL/min. LC conditions were set at 90 % (A) ramped linearly over 8.0 min to 100% (B) and held until 10.0 min at 100% B. At 10.0 min the gradient was switched back to 90% (A). GW, YK, MN, and 7-methoxy-MN were synthesized by the published methods[1].

**MN-Kinobead.** To a stirred solution of 7-methoxy-MN (352 mg, 1.101 mmol, 1.0 equiv.) in dry dichloromethane (10 mL), boron tribromide ($BBr_3$) (1 M solution of $BBr_3$ in DCM, 2.7 mL, 2.753 mmol, 2.5 equiv.) was added at 0 °C, and the reaction mixture was stirred at room temperature under a nitrogen atmosphere for 16 hours. Then, boron tribromide (1 M solution of $BBr_3$ in DCM, 2.7 mL, 2.753 mmol, 2.5 equiv.) was added again at 0 °C and stirred at room temperature for another 16 hours. The reaction progress was monitored by TLC and LCMS. After the reaction was completed, the mixture was cooled with an ice bath, and aqueous sodium bicarbonate (30 mL) was added and stirred for 5 minutes. Dichloromethane (20 mL) was then added, and the mixture was extracted with dichloromethane (2 x 20 mL) and dried over sodium sulfate. The organic layer was evaporated to give a crude mass, which was purified by flash column chromatography (0 to 15% gradient elution of MeOH in DCM) to give 7-hydroxy-MN (150 mg, 47% yield): $^1$H NMR (850 MHz, DMSO-$d_6$) δ 8.74 – 8.67 (m, 2H), 8.11 (dd, J = 7.5, 0.7 Hz, 1H), 7.59 – 7.51 (m, 2H), 7.51 – 7.43 (m, 2H), 7.21 – 7.14 (m, 2H), 7.08 (d, J = 2.6 Hz, 1H), 6.64 (dd, J = 7.5, 2.6 Hz, 1H), 3.87 (s, 3H); $^{13}$C NMR (214 MHz, DMSO-$d_6$) δ 161.73 (d, J = 245.1 Hz) 158.20, 150.76, 146.46, 141.50, 137.15, 130.43, 130.41, 129.71, 129.67, 124.77, 124.63, 116.96, 115.44, 115.34, 107.59, 94.67, 55.79; $^{19}$F NMR (376 MHz, $CDCl_3$) δ -113.48; HRMS calculated for $C_{18}H_{13}FN_3O$ $[M+H]^+$: 306.1042. Found: 306.1033.

To a solution of 7-hydroxy-MN (411 mg, 1.35 mmol) in DMF (5.0 mL) was added 2-((tert-butoxycarbonyl)amino)ethyl 4-methylbenzenesulfonate (849 mg, 2 eq, 2.69 mmol) and cesium carbonate (1.32 g, 3 eq, 4.04 mmol). The mixture was stirred for 2 h at 80 °C, turning a deep red. The solvent was blown down overnight and the resulting red gum was dissolved in 40% TFA in DCM (5.0 mL). The mixture was stirred at room temperature overnight. The solvent was blown down and crude product was purified by preparative HPLC (column: Phenomenex, Luna 5 μM Phenyl-Hexyl, 100 Å, 75 × 30 mm, 5 micron; mobile phase A: water (0.05% TFA), B: methanol; method: 20–100% B, 6 min + 100% B, 4 min). 7-(2-aminoethoxy)-MN was obtained as a bright yellow crystalline solid (161 mg, 23%): $^1$H NMR (400 MHz, DMSO) δ 8.86 – 8.80 (m, 2H), 8.40 (d, *J* = 7.6 Hz, 1H), 8.19 (s, 2H), 7.72 – 7.65 (m, 2H), 7.62 – 7.52 (m, 2H), 7.36 (d, *J* = 2.5 Hz, 1H), 7.34 – 7.24 (m, 2H), 6.91 (dd, *J* = 7.6, 2.5 Hz, 1H), 4.39 (t, *J* = 4.9 Hz, 2H), 3.33 (t, *J* = 5.2 Hz, 2H); $^{13}$C NMR (100 MHz, DMSO) δ 149, 131, 131, 127, 126, 117, 116, 110, 95, 66, 38; HRMS calculated for $C_{38}H_{34}BF_3N_7O3$ $[M+H]^+$: 704.2768. Found: 704.2756.

7-(2-aminoethoxy)-MN (62 mg) was placed into a 20 mL scintillation vial and dissolved in 8 mL of 50:50 (v:v) DMF/EtOH. 8 mL of a slurry of ECH-Sepharose 4B in 20% ethanol was distributed equally between 2 Bio-Rad Poly-Prep Columns (Product # 731-1550). The ethanol was left to drain, followed by washing with equal volume 50:50 DMF/EtOH. Columns were capped at the bottom and 4 mL of 7-(2-aminoethoxy)-MN solution was pipetted equally into each column. of EDC•HCl (153.6 mg, Sigma: Cat#: E1769) was added to each column, followed by capping the top and agitating for solubilizing and bead resuspension. The columns were wrapped in aluminum foil and left to tumble at 4 °C overnight. The columns were allowed to warm to room temperature. Liquid was drained by gravity flow, the column was recapped at the bottom, and 1M ethanolamine in 50:50 DMF/EtOH (4 mL) and EDC.HCl (153.6 mg) were added to block any unreacted sites. The columns were capped, agitated, wrapped in aluminum foil, and tumbled at room temperature for 2 h. After mixing, liquid was drained and the columns were washed with 8 mL each: 50:50 DMF/EtOH, 2X acetate wash (acetic acid, NaCl, pH 4.0, filtered), 2X Tris wash (Trizma, NaCl, pH 8.3, filtered), 2X acetate wash, 2X Tris wash, and 20% ethanol. Columns were capped at the bottom and MN-kinobeads resuspended in equal volume bead to 20% ethanol to be placed in 15 mL Falcon tubes. The MN-kinobeads were stored in a 4 °C refrigerator.

**MN-Bodipy.** To a stirred solution of 7-(2-aminoethoxy)-MN (40 mg, 0.13 mmol, 1.0 equiv.) in DMF (4 mL), 2-(2-((tert-butoxycarbonyl)amino)ethoxy)ethyl 4-methylbenzenesulfonate (47 mg, 0.13 mmol, 1.0 equiv.) and $K_2CO_3$ (72 mg, 0.52 mmol, 4.0 equiv.) were added at rt, and the reaction mixture was stirred at 50 °C for 16 h. After the reaction was complete by TLC and LCMS, water (25 mL) was added, the mixture was extracted with ethyl acetate (2 x 25 mL) and dried over sodium sulfate. The organic layer was evaporated to give a crude product, which was purified by flash column chromatography (0–5% gradient elution of MeOH in DCM) to give the Boc-amino-PEG-MN product as a pale yellow liquid (27 mg, 42% yield): $^{1}$H NMR (400 MHz, $CDCl_3$) δ 8.74 (s, 2H), 7.93 (d, $J$ = 7.6 Hz, 1H), 7.60 – 7.50 (m, 2H), 7.38 – 7.31 (m, 2H), 7.08 – 6.95 (m, 3H), 6.64 (dd, $J$ = 7.5, 2.4 Hz, 1H), 5.00 (broad s, 1H), 4.24 – 4.17 (m, 2H), 3.91 – 3.82 (m, 2H), 3.61 (t, $J$ = 5.2 Hz, 2H), 3.37 – 3.30 (m, 2H), 1.42 (s, 9H); $^{13}$C NMR (126 MHz, $CDCl_3$) δ162.84 (d, $J$=248.19 Hz), 158.21, 156.08, 151.12, 146.85, 142.25, 137.66, 130.21, 130.14, 129.15, 129.12, 124.52, 123.76, 117.20, 115.81, 115.63, 108.78, 95.57, 79.46, 70.69, 69.04, 68.06, 40.40, 28.51; $^{19}$F NMR (376 MHz, $CDCl_3$) δ -113.10; HRMS calculated for $C_{22}H_{22}FN_4O_2$ $[M-(Boc+H_2)+H]^+$: 393.1727. Found: 393.1716.

Boc-amino-PEG-MN was dissolved in 20% TFA in DCM and stirred at room temperature for 30 min. TLC confirmed the disappearance of the starting material. The solvent was removed under reduced pressure, and the resulting crude pale yellow liquid was dissolved in DMF and BODIPY NHS ester (7.9 mg, 1 eq, 18 µmol) and DIPEA (12 mg, 16 µL, 5 eq, 92 µmol) were added. The mixture was stirred for 30 min and then subjected to preparative HPLC (column: Phenomenex, Luna 5 µM Phenyl-Hexyl, 100 Å, 75 × 30 mm, 5 micron; mobile phase A: water (0.05% TFA), B: methanol; method: 20–100% B, 6 min + 100% B, 4 min). The desired tracer MN-Bodipy was obtained as a dark purple solid (6.4 mg, 32%): $^{1}$H NMR (500 MHz, MeOD) δ 8.68 – 8.64 (m, 2H), 8.27 (d, $J$ = 7.6 Hz, 1H), 7.47 – 7.42 (m, 2H), 7.40 – 7.32 (m, 2H), 7.20 – 7.15 (m, 4H), 7.14 (s, 1H), 7.13 – 7.08 (m, 3H), 6.93 (d, $J$ = 4.6 Hz, 1H), 6.87 (d, $J$ = 3.9 Hz, 1H), 6.37 – 6.28 (m, 2H), 4.39 – 4.32 (m, 2H), 3.94 – 3.88 (m, 2H), 3.68 (dd, $J$ = 5.6, 4.7 Hz, 2H), 3.46 (t, $J$ = 5.1 Hz, 2H), 3.15 (t, $J$ = 7.5 Hz, 2H), 2.57 (t, $J$ = 7.7 Hz, 2H); $^{13}$C NMR (126 MHz, MeOD) δ 174.98, 165.29 (d, $J$ = 250.74 Hz), 165.09, 156.20, 152.01, 150.73, 144.25, 138.81, 136.90, 134.78, 133.23, 132.08, 132.01, 128.27, 127.44, 127.14, 126.58, 124.71, 124.47, 124.06, 121.06, 119.81, 119.24, 117.62, 117.44,

117.24, 112.74, 112.46, 93.26, 70.88, 70.64, 69.78, 40.24, 36.02, 25.91; $^{19}$F NMR (470 MHz, MeOD) δ -77.35; HRMS calculated for $C_{38}H_{34}BF_3N_7O_3$ [M + H]+: 704.2768. Found: 704.2756.

**CK1 inhibitors.** GW271431X, GSK1660450B, and GSK507358A were available from the kinase inhibitor collection within the SGC-UNC with a purity > 98%. CDK1/2 Inhibitor III was obtained from Millipore Sigma and all other inhibitors were obtained from MedChemExpress as either solid or 10mM DMSO stock solution with purities ≥ 98%. All solid samples used were made into 10mM DMSO stock solutions and stored at -20 °C.

**HEK293 Cell Lysate.** Expi293 cells (Thermo; a phenotype of HEK293F selected for high density growth and robust protein expression) were grown to approximately $3.0x10^6$ cells/mL (log phase growth, well below saturation) in Expi293 medium following standard suspension tissue culture protocols from the manufacturer. The cells were collected by centrifugation at 300xg at 4 °C for 20 minutes. The pellets were gently washed with ice cold PBS. The collection and washing process was repeated two more times. Then the pellets were resuspended in cold PBS and portioned into $100–150x10^6$ cell aliquots. Each aliquot was pelleted one last time, the PBS siphoned off, and the pellet frozen at -80 °C until processing. Each aliquot of $100–150x10^6$ cells gave roughly 5 mg total protein in processing. Cell pellets were thawed on ice and introduced to fresh MIB lysis buffer (50 mM HEPES, 150 mM NaCl, 0.5% Triton X-100, 1mM EDTA, 1mM EGTA, 10 mM NaF, 2.5 mM $NaVO_4$, pH 7.5, Phosphatase Inhibitor Cocktail 2 & 3 [Sigma P5726 and P0044], cOmplete Protease Inhibitor Cocktail [EDTA free, Roche). Cells were incubated in MIB lysis buffer for 10 min on ice with tube inverting every 2 min. Cells were sonicated using a Qsonica ultrasonicator at 125 W, 20 kHz, 35% amplitude, in 3x10 sec treatments, allowing sample to cool on ice for 30 sec between pulses. Cell lysate was clarified through 0.20 μm syringe filters (Corning, 431219) into a single 15 mL prechilled Falcon tube. Protein concentration was determined by Bradford method.

**Fungal Strain and Culture Conditions.** Archives of all strains were maintained at -80 ˚C in 25% glycerol. Strains were grown in standard conditions at 30 ˚C in YPD (1% yeast extract, 2% peptone, 2% dextrose), unless otherwise indicated. Active cultures were maintained on solid YPD (2% agar) at room temperature no longer than 5 days. Fungal strains include: CaLC2742[2], CaLC249[2], CaSS1 *tetO-YCK2/yck2*Δ[3], and CaSS1 *tetO-ERG6/erg6*Δ[4].

***C. albicans*** **Cell Lysate.** Archives of all strains were maintained at -80 °C in 25% glycerol. Strains were grown in standard conditions at 30 °C in YPD (1% yeast extract, 2% peptone, 2% dextrose), unless otherwise indicated in RPMI (10.4 g/L RPMI powder with L-glutamine (Gibco), 165 mM MOPS, 2% glucose, 5 mg/mL histidine, pH 7), or SD (2% glucose, 6.7 g/L yeast nitrogen base without amino acids).

*C. albicans* strain SC5315 was grown to mid-late log phase (OD 3–4) in four liters YPD medium at 30 °C and harvested by centrifugation at 3,000 g. The pellet was transferred into a 60 mL syringe and squeezed into liquid nitrogen. Cells were disrupted by grinding in liquid nitrogen using mortar-pestle. Each gram of the resulting powder was dissolved in 2 mL of 1.5x MIB buffer (75 mM HEPES-NaOH pH 7.5, 225 mM NaCl, 0.75% Triton X-100, 1.5 mM EDTA, 1.5 mM EGTA, 15 mM NaF, 3.75 mM Na3VO4, 7.5%(v/v) glycerol) supplemented with cOmplete proteinase inhibitor (Roche; 2x of the recommended concentration) and phosphatase inhibitor cocktail II (Sigma). The lysate was sonicated for 20 min (10 sec on; 10 sec off) at 30% amplitude using a Misonix S-4000 dual horn sonicator with 3/4-inch probes and spun at 17,000 rpm in a JA25.50 rotor for 10 min. The supernatant was further clarified by a 90 min centrifugation in a Ti45 rotor at 40,000 rpm. Protein concentration was determined by Bradford method using a 1:10 diluted lysate in 1x TE buffer (10 mM Tris-HCl pH 7.5, 1 mM EDTA).

**Western Blot.** A volume of 50 μL of resuspended kinobeads was added to a 1.5 mL Eppendorf tube containing 1 mg total protein from HEK293 cell lysate. Volume was brought to 1 mL by adding 1X TBST (Fischer, Lot# 235915). Samples were placed on a tumbler and left to mix at 4 °C overnight. The samples were centrifuged at 5000 rpm for 1 min at room temperature, later pipetting out all supernatant. Samples were washed with 1 mL of 1X TBST, vortexed, centrifuged as before, and supernatant removed 4 times. 40 μL of 1X Tris-Glycine SDS Sample Buffer (Novex Cat# LC2676) was added, followed by heating tubes in a heating block at 85 °C for 5 min. Samples were spun down as before, transferring 40 μL supernatant to another Eppendorf tube. Untreated cell lysate sample was prepared similarly in 40 μL with 1 ug protein, 1X NuPAGE Sample Reducing Agent (Invitrogen, Cat# 0004), 1X TBST, and 1X Tris-Glycine SDS Sample Buffer. Cell lysate sample was heated and centrifuged as previously stated for samples containing kinobeads. 1X Tris-Glycine SDS gel running buffer (Novex, Cat# LC2675) was added to the running tank of and XCell SureLock Mini-Cell. Novex 4–12% Tris-Glycine Plus WedgeWell Gel (Invitrogen, Cat# XP04120BOX) was loaded into the running tank, followed by pipetting 5 μL ladder (Invitrogen, REF LC5925) and 40 μL samples or 1X Tris-Glycine SDS into unused wells. The gel was run at 200V for 40 min. The gel was removed from the plastic tray and placed into the iBlot Gel Transfer Device on top of the anode stack. Pre-soaked iBlot filter paper was placed on top of the gel, bubbles rolled out, and the cathode stack placed on top of the filter paper. Blotting took place using protocol P3 for 10 min. The membrane was rocked on a rocker in 1X TBST + 5% w/v milk as blocking buffer for 30 min at room temperature. The membrane was washed, then transferred to the Invitrogen Bandmate Automated Western Blot Processor using the following procedure: blocked for 1 h at rt in 5% milk in TBST, incubation with primary antibody p38 MAPK (p38-3f11, Thermo Fisher Cat# 33-1300, 1:1000) for 2 h at rt in 5% milk in TBST, followed by

Rb-HRP (1:5000) in TBST at rt for 1h. SuperSignal West Femto reagent (Thermo Fisher,Cat# 34094) was used for imaging on Invitrogen iBright Imager.

**MIB/MS Assay.** Protein lysate containing ≥ 5 mg total protein in 4.0 mL MIB lysis buffer was mixed with 4 μL DMSO or test compound followed by 1 h incubation on ice. Bio-Rad Poly-Prep Columns (Product # 731-1550) were prepared for each sample with 350 μL of a 50% slurry of a total MIB matrix mix; kinobead matrix consists of Shokat, PP58, Purvalanol B, UNC-21474 (14% each by volume), VI-16832, and Ctx-0294885 (22% each by volume) on ECH Sepharose 4B in 20% aq. ethanol. Columns were washed with 2 mL high salt MIB wash buffer (50 mM HPES, 1M NaCl, 0.5% Triton X-100, 1 mM EDTA, 1 mM EGTA, pH 7.5) for initial equilibration. Cell lysate samples were brought to 1M NaCl. Lysate samples were pipetted onto columns, allowing flow through to pass. Columns were flushed with 5 mL high salt MIB wash buffer, low salt MIB wash buffer (50 mM HPES, 150 mM NaCl, 0.5% Triton X-100, 1 mM EDTA, 1 mM EGTA, pH 7.5), and 500 μL 0.1% SDS in low salt MIB wash buffer. For MIB extraction, the columns were capped and the kinobeads were resuspended with 500 μL MIB elution buffer (0.5% w/v SDS, 0.1 M Tris-HCl, pH 7.25) followed by heating at 95 °C for 10 min. This step was repeated one more time. The proteins released from the MIB were reduced with 5mM DL-Dithiothreitol in a heating block for 25 min at 60 °C. Cysteine alkylation was performed with 18 mM iodoacetamide in a dark chamber at room temperature for 30 min. After dark incubation, alkylation was quenched by adding more DL-Dithiothreitol for a final concentration of 10 mM protein concentration was performed with Amicon Millipore Ultra-4 10K cutoff spin columns (Cat# UFC801008). Samples in the 10 kDa filters were centrifuged for 30 min at 3000 rpm, 4 °C. Methanol/chloroform extraction was performed on the concentrated samples. Briefly, 400 μL methanol, 100 μL chloroform, and 300 μL LC-MS grade water was pipetted into each Eppendorf tube. Samples were vortexed for 10 sec and centrifuged at 4 °C for 10 min at 15,000 rpm, forming an interphase. The upper aqueous layer was pipetted out and the extracts were washed four times with 500 μL methanol. Samples were dried down on a Labconco Acid-Resistant CentriVap Concentrator for 30 min, followed by reconstitution in 100 uL of 50 mM HEPES, pH 8.0 buffer in LC-MS grade water. 3 μL of 0.4 μg/μL sequencing-grade Modified Trypsin (Promega V5111) was pipetted into each sample, mixed, and incubated at 37 °C overnight for digestion.

On day 3, trypsin-digested samples were washed 3 times with ethyl acetate, discarding the top layer after each rinse. Samples were then dried down on the Labconco Acid-Resistant CentriVap Concentrator for 2 h. Dried trypsin-digested samples were desalted using C-18 PepClean Spin Columns (Pierce, Cat# 89870). Dried pellets were resuspended in 200 μL equilibration buffer (5% MeCN, 0.5% TFA in LC-MS water) and loaded onto the wetted columns. Columns were washed four times with 200 μL 5% MeCN/0.5% TFA in LC-MS grade water, then two times with 25 μL 50% MeCN/0.5% TFA in LC-MS grade water.

Samples were dried down on the Labconco Acid-Resistant CentriVap Concentrator for 3 h and stored at -80 °C. For LC-MS/MS analysis, the dried tryptic peptides were resuspended in 15 uL 2% Acetonitrile, 0.1% Formic Acid. LC-MS/MS analysis was performed with a Thermo Scientific Easy nLC 1200 coupled to a Thermo Scientific Biopharma QExactive HF Orbitrap Mass Spectrometer equipped with an Easy-Spray Nano Source. Tryptic peptides were chromatographically separated using an Easy-Spray PepMap C18 column (75 µm ID X 25cm, 2 µm particle size; Thermo Scientific) and eluted over a 90 min method. Separation was achieved with a gradient of 5–45% B at a 250 nl/min flow rate with mobile phase A (water with 0.1% formic acid, v/v) and mobile phase B (80% acetonitrile, 0.1% formic acid, v/v). The QExactive HF was operated in data-dependent mode where the 15 most intense precursors were selected for subsequent fragmentation. Resolution for the precursor scan (m/z 375–1700) was set to 60,000 with an AGC target of $3\times10^6$ ions, 100 ms max IT. MS/MS scans resolution was set to 15,000 with an AGC target value of $1\times10^5$ ions, 100 ms max IT. The normalized collision energy was set to 27% for HCD. Dynamic exclusion was set to 30 sec and precursors with unknown charge or a charge state of 1 and $\geq$ 7 were excluded. The data was searched with MaxQuant (version 1.6.15.0) against a reviewed human database (containing 20404 entries, downloaded January 2024) or against *C. albicans* SC5314 database (containing 6,039 entries, downloaded July 2024) and a contaminants database using Andromeda within MaxQuant. Enzyme specificity was set to trypsin, allowing for two missed cleavages. Methionine oxidation and N-terminal acetylation were set as dynamic modifications. Carbamidomethylation on cysteine residues was set as fixed modification. Match between runs was enabled with a 4 min and 20 min matching and alignment time windows, respectively. All *C. albicans* proteins and human kinases were imported into Perseus (version 1.6.14.0) for further processing. Within Perseus, decoy proteins, contaminants, single hits, and proteins with 50% missing values were removed from analysis. $Log_2$ transformation of LFQ intensity was performed. $Log_2$ fold change (FC) ratios were calculated using the $Log_2$ LFQ intensities of drug treated sample compared to control.

**Global Proteomic Analysis.** Samples were stored in MIB lysis buffer before LC-MS analysis. For each sample 100 µg of protein lysate was acetone precipitated then dried down and resuspended in 100 µl of 1M urea, 50 mM ammonium bicarbonate, pH 8. Next, samples were reduced with dithiothreitol, alkylated with iodoacetamide, digested with LysC (Wako) at 37 °C for 2 h at a 1:25 enzyme:protein ratio, and trypsin (Promega) overnight at 37 °C at a 1:25 enzyme:protein ratio. The resulting peptides were acidified to 0.5% trifluoroacetic acid (TFA; Pierce), dried, and desalted using Thermo desalting spin columns. Eluates were dried by vacuum centrifugation and peptide concentration was determined by Pierce Quantitative Fluorometric Assay and all samples were normalized to 0.1 µg/µl and subjected to LC-MS/MS analysis using a VanquishNeo coupled to an Orbitrap Astral mass spectrometer (Thermo Scientific). Samples were injected onto an IonOptics Aurora series 3 C18 column (75 µm id × 15 cm, 1.6 µm particle size; IonOpticks) and separated over a 60 min method. The gradient for separation consisted of 2–30% mobile phase B at a 300 nl/min flow rate, where mobile phase A was 0.1% formic acid in water and mobile phase B consisted of 0.1% formic acid in 80% ACN. Astral was operated in product ion scan mode for Data Independent Acquisition (DIA). A full MS scan (m/z 380–980) was collected; resolution was set to 240,000 with a maximum injection time of 5 ms and AGC target of 500%. Following the full MS scan, a product ion scan was collected (30,000 resolution); AGC target set to 500%; maximum injection time set to 3 ms; precursor mass range set to 380–980 m/z; the isolation window was set to 3 m/z. Raw data files were processed using Spectronaut (version 18.7.240506; Biognosys) and searched against the *C. albicans* strain SC5314 database (UP000000559, containing 6,039 entries, downloaded May 2024) appended with a common contaminants database (245 sequences) using the Pulsar search algorithm. Enzyme specificity was set to trypsin, up to two missed cleavage sites were allowed; methionine oxidation, and Protein N-term acetylation were set as variable modifications and carbamidomethylation of Cys was set as a fixed modification. Data was normalized using the global normalization strategy and no imputation was performed. All data were filtered at a 1% false discovery rate (FDR), and proteins were filtered for a minimum of 1 peptide.

**MN-Kinobead Pulldown.** The protein pulldown experiment was conducted using a modified MIB/MS protocol with 350 µL of the MN-kinobead. Samples were prepared as biological triplicates and introduced to DMSO or MN (10 µM) during the experiment. The resulting dried tryptic peptides were resuspended in 15 µL 2% acetonitrile, 0.1% formic acid. The peptide samples were analyzed by LC-MS/MS using an Ultimate3000 coupled to an Exploris480 mass spectrometer (Thermo Scientific). Samples were injected onto an IonOpticks Aurora series 3 C18 column (75 µm id × 15 cm, 1.6 µm particle size; IonOpticks) and separated using a 90 min method. The gradient for separation consisted of 2–40% mobile phase B at a 250 nl/min flow rate, where mobile phase A was 0.1% formic acid in water and mobile phase B consisted of 0.1% formic acid in ACN. The Exploris480 was operated in data-dependent mode with a cycle time of 2s. Resolution for the precursor scan (m/z 375–1500) was set to 120,000, while MS/MS scans resolution was set to 15,000. The normalized collision energy was set to 30% for HCD. Peptide match was set to preferred, and precursors with unknown charge or a charge state of 1 and ≥ 7 were excluded. Raw data files were processed using Proteome Discoverer version 3.1 and searched against the *C. albicans* SC5314 database (containing 6,039 entries, downloaded July 2024) appended with a common contaminants database (245 sequences) using the Sequest HT search algorithm. Enzyme specificity was set to trypsin, up to two missed cleavage sites were allowed; methionine oxidation, and carbamidomethylating of Cys were set as variable modifications. A precursor mass tolerance of 10ppm and fragment mass tolerance of 0.02 Da were used. Label-free quantification (LFQ) was enabled. Data were filtered based on a 1%/5% peptide/protein false discovery rate (FDR), a minimum of 2 peptides, and quantitation in minimum 3 samples. Data was not normalized. Further processing and Statistical analysis were conducted in Perseus (version 1.6.14.0). Missing values were imputed from a normal distribution. Student's t-test was performed for the MN to DMSO comparison and a p-value $< 0.05$ was considered statistically significant. LFQ $log_2$ fold change ratios were calculated, and an absolute $log_2$ ratio of 1 or greater was considered significant.

**Biochemical Assay.** Kinase assays were performed using the ADP-Glo kinase assay kit (Promega) in solid white 384 well plates (Corning). Assays with were performed in kinase buffer (1x: 2 mM NaHEPES pH 7.5, 650 mM KCl, 50 mM $MgCl_2$, 25 mM β-glycerophosphate) supplemented with 2 µg casein kinase 1 peptide substrate (SignalChem) per reaction, 0.05 µg pure p38α (Promega V2701), ~0.115 µg purified recombinant Yck2 kinase domain, and 0.05 µg purified CK1α (Abcam) per reaction as indicated. Each kinase inhibitor of interest was added in a two-fold dilution series at the concentrations indicated, followed by the addition of ATP at 20 µM (Yck2 $K_m$ATP), 10 µM (CK1α) or 80 µM (p38α), respectively. Assays were performed in 10 µL reactions ($n = 3$) and incubated for 30 min at 30 °C. ADP-Glo kinase assay reagents (Promega) were applied to assay wells per manufacturer's instructions. Luminescence was measured with a TECAN Spark multimode microplate reader, and background luminescence was subtracted from reaction wells from control wells incubated with all reaction reagents except for the relevant kinase enzymes. $IC_{50}$ values were calculated, and data plotted using GraphPad Prism 9's Nonlinear fit function.

**Anti-fungal Assay.** All tested compounds were dissolved in DMSO. Drug susceptibility assays were performed in 384-well plates in a final volume of 0.04 mL/well (~1 x $10^3$ cells/mL *C. albicans*) with two-fold dilutions of each compound in YPD or RPMI medium (10.4 g/L RPMI powder with L-glutamine (Gibco), 165 mM MOPS, 2% glucose, 5 mg/mL histidine, pH 7), as indicated. Plates were incubated in the dark at 30 ˚C or 37 ˚C with 5% $CO_2$ under static conditions. Absorbance was measured at 600 nm after the indicated incubation times using a spectrophotometer (Molecular Devices) to assess growth. Growth was normalized to the no-drug controls. All assays represent average values of technical duplicates, and were performed with two biological replicates to confirm reproducibility. Data was quantitatively displayed as heat maps using the program Java TreeView3. For assays utilizing the conditional expression strains from the Gene Replacement and Conditional Expression (GRACE) collection, strains were grown overnight in the presence and absence of the indicated doxycycline concentration(s) to repress gene expression.

**NanoBRET Assay.** Synthetic DNA encoding NanoLuc and a linker sequence of GSSGAIA was fused to the N-terminus of the kinase domain of Yck2 (residues S37–N345) and cloned into the pTwist CMV Puro plasmid (Twist Biosciences) to generate NLuc-Yck2.[5] HEK293 cells in a 96-well plate expressing the

NLuc-Yck2 fusion protein were mixed with an increasing concentration of MN-Bodipy tracer. The resulting BRET ratio was fitted to the sigmoidal 3-parameter dose–response curve using Prism 9.0 (GraphPad, Boston MA, USA). As a control, the same experiment was repeated in the presence of an excess of unlabeled compound MN (20 μM) as a competitive inhibitor for 2 h before adding 3X Complete Substrate plus Inhibitor Solution. MN-Bodipy at 500 nM was selected as optimal tracer concentration for the Yck2 NanoBRET assay.

NanoBRET Assays were run with a modified version of the previously published protocols[6,7]. HEK293 cells were cultured at 37 °C in 5% $CO_2$ in Dulbecco's modified Eagle medium (DMEM; Gibco) supplemented with 10% fetal bovine serum (VWR/Avantor). A transfection complex of DNA at 10 μg/mL was created, consisting of 9 μg/mL carrier DNA (Promega) and 1 μg/mL NLuc-kinase fusion DNA in Opti-MEM without serum (Gibco). FuGENE HD (Promega) was added at 30 μL/mL to form a lipid:DNA complex. The solution was then mixed and incubated at room temperature for 20 min. The transfection complex was mixed with a 20× volume of HEK293 cells in DMEM/FBS to arrive at a final concentration of 200,000 cells/mL, and 100 μL/well was added to a 96-well plate that was incubated overnight at 37°C and 5% $CO_2$. The following day, the media were removed via aspiration and replaced with 85 μL of Opti-MEM without phenol red. A total of 5 μL per well of 20× NanoBRET Tracer was added. Thus final tracer concentrations were as follows; Yck2:MN-Bodipy 500nM, CSNK1A1:Promega K8 300 nM, CSNK1D:Promega K10 at 500 nM, CSNK1E:Promega K8 at 500 nM, CSNK1G2:Promega K8 at 130 nM, and p38α:Promega K4 at 31 nM in Tracer Dilution Buffer (Promega N291B) was added to all wells, except the "no tracer" control wells. Test compounds (10 mM in DMSO) were diluted 100× in Opti-MEM media to prepare stock solutions and evaluated at 11 concentrations. A total of 10 μL per well of the 10-fold test compound stock solutions (final assay concentration of 0.1% DMSO) were added. For "no compound" and "no tracer" control wells, DMSO in Opti-MEM was added for a final concentration of 1.1% across all wells; 96-well plates containing cells with NanoBRET Tracer and test compounds (100 μL total volume per well) were equilibrated (37°C/5% $CO_2$) for 2 h. The plates were cooled to room temperature for 15 min. The NanoBRET NanoGlo substrate (Promega) at a ratio of 1:166 to Opti-MEM media in combination with an extracellular NLuc Inhibitor (Promega) diluted at 1:500 (10 μL of 30 mM stock per 5 mL of the Opti-MEM plus substrate) was combined to create a 3× stock solution. A total of 50 μL of the 3× substrate/extracellular NL inhibitor was added to each well. The plates were read within 30 min on a GloMax Discover luminometer (Promega) equipped with a 450 nm BP filter (donor) and 600 nm LP filter (acceptor) using 0.3 s of integration time. Raw milliBRET (mBRET) values were obtained by dividing the acceptor emission values (600 nm) by the donor emission values (450 nm) and multiplying by 1000. Averaged control values were used to represent complete inhibition (no tracer control: Opti-MEM + DMSO only) and no inhibition (tracer only control: no compound, Opti-MEM + DMSO + Tracer only) and were plotted alongside the raw mBRET values. The data was first normalized and then fit using the Sigmoidal 3PL binding curve in Prism Software to determine $IC_{50}$ values.

The K192 NanoBRET selectivity assay was run according to the Draft Promega technical manual. Reagents were supplied by Promega (Promega NP 4101). For the assay, DNA from the prepared kinase vector panel plates A&B were mixed with Fugene in 96 well plates (Corning 3917) and incubated at room temperature for 30 minutes. Control vectors used are the NanoLuc Low control vector is pNL1.1.CMV [Nluc/CMV] Vector (Cat.# N1091) and the transfection control vector is NanoLuc-HIPK2 Fusion Vector (Cat.# NV3221). HEK293 cells were grown to 75–95% confluency in DMEM (Gibco 11995-065) supplemented with FBS (Avantor 97068-085) at 37 °C in 5% RH. On the first day of the assay, cells were harvested and resuspended in Opti-MEM (Gibco 11058-021) supplemented with 1% FBS (Avantor 97068-085) at $2.5 \times 10^5$ cells per mL. 60 µL of cell suspension was mixed with 10 µL of prepared DNA (10X concentration) and 30 µL of Fugene (30 µL/mL in Opti-MEM) as outlined by Promega and incubated overnight at 37°C in a 5% $CO_2$ incubator. On day two of the assay, 5 µL of 20X Promega K10 tracer was prepared and added at the recommended concentrations. Next, 10 µL of test inhibitor YK or MN at 1 µM in Opti-MEM (diluted from a 10 mM solution in DMSO) was added to the test wells while an equivalent volume of Opti-MEM was added to the high-control wells. Plates were kept at 37 °C in a 5% $CO_2$ incubator for two hours. After 2 h, plates were allowed to equilibrate to room temperature for 15 min. A solution of 3X Complete Substrate plus Inhibitor Solution was freshly prepared, consisting of a 1:166 dilution of NanoBRET Nano-Glo Substrate plus a 1:500 dilution of Extracellular NanoLuc Inhibitor in Opti-MEM medium without serum or phenol red. 50 µL of the 3X Complete Substrate plus Inhibitor Solution was added to each assay well, including control wells. After 2–3 minutes, the plate was shaken at 300 rpm for 10 sec and the donor emission wavelength (450 nm) and acceptor emission wavelength (610 nm) were measured using the Glomax Discover System. As a quality check, the donor signal-to-background ratio was calculated for each individual kinase by dividing the mean donor signal for each kinase by the mean donor signal for the signal-to-background control wells. Fractional occupancy for the test drug for each kinase was determined using the following formula:

$$\text{Occupancy (\%)} = [\ 1 - (\text{Sample} - \text{Bottom}) / (\text{Top} - \text{Bottom})\ ] \times 100$$

Where:

Sample = Mean BRET value across all Sample (tracer + compound) wells for an individual kinase.

Top = Mean BRET value across all Top (tracer + vehicle) control wells for an individual kinase.

Bottom = Mean BRET value of NanoLuc control wells.

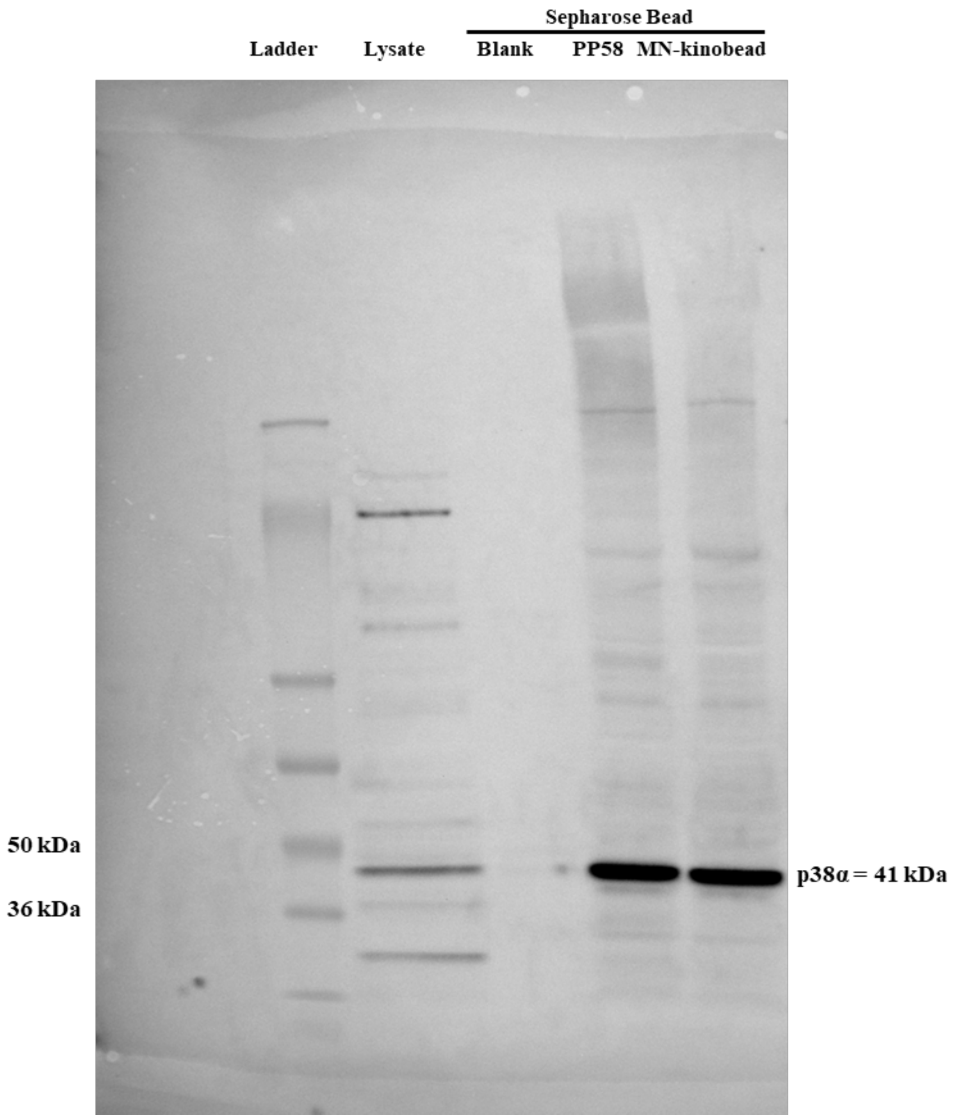


**Figure S1.** Western-blot for kinobead pulldown of p38α kinase from HEK293 cell lysate.

### References


(1) Puumala, E.; Nandakumar, M.; Williams, N. S.; Stogios, P. J.; Zarnowski, R.; Liu, Z.; Whitesell, L.; Robbins, N.; Andes, D.; Willson, T. M.; Cowen, L. E. Structure-Guided Optimization of Small Molecules Targeting the Yeast Casein Kinase, Yck2, as a Therapeutic Strategy to Combat Candida Albicans. **2024**.

(2) Calabrese, D.; Bille, J.; Sanglard, D. A Novel Multidrug Efflux Transporter Gene of the Major Facilitator Superfamily from Candida Albicans (FLU1) Conferring Resistance to Fluconazole. *Microbiology (N Y)* **2000**, *146* (11), 2743–2754. https://doi.org/10.1099/00221287-146-11-2743.

(3) Fu, C.; Zhang, X.; Veri, A. O.; Iyer, K. R.; Lash, E.; Xue, A.; Yan, H.; Revie, N. M.; Wong, C.; Lin, Z.-Y.; Polvi, E. J.; Liston, S. D.; VanderSluis, B.; Hou, J.; Yashiroda, Y.; Gingras, A.-C.; Boone, C.; O'Meara, T. R.; O'Meara, M. J.; Noble, S.; Robbins, N.; Myers, C. L.; Cowen, L. E. Leveraging Machine Learning Essentiality Predictions and Chemogenomic Interactions to Identify Antifungal Targets. *Nat Commun* **2021**, *12* (1), 6497. https://doi.org/10.1038/s41467-021-26850-3.

(4) Roemer, T.; Jiang, B.; Davison, J.; Ketela, T.; Veillette, K.; Breton, A.; Tandia, F.; Linteau, A.; Sillaots, S.; Marta, C.; Martel, N.; Veronneau, S.; Lemieux, S.; Kauffman, S.; Becker, J.; Storms, R.; Boone, C.; Bussey, H. Large-scale Essential Gene Identification in *Candida Albicans* and Applications to Antifungal Drug Discovery. *Mol Microbiol* **2003**, *50* (1), 167–181. https://doi.org/10.1046/j.1365-2958.2003.03697.x.

(5) Robers, M. B.; Dart, M. L.; Woodroofe, C. C.; Zimprich, C. A.; Kirkland, T. A.; Machleidt, T.; Kupcho, K. R.; Levin, S.; Hartnett, J. R.; Zimmerman, K.; Niles, A. L.; Ohana, R. F.; Daniels, D. L.; Slater, M.; Wood, M. G.; Cong, M.; Cheng, Y.-Q.; Wood, K. V. Target Engagement and Drug Residence Time Can Be Observed in Living Cells with BRET. *Nat Commun* **2015**, *6* (1), 10091. https://doi.org/10.1038/ncomms10091.

(6) Elkins, J. M.; Fedele, V.; Szklarz, M.; Abdul Azeez, K. R.; Salah, E.; Mikolajczyk, J.; Romanov, S.; Sepetov, N.; Huang, X.-P.; Roth, B. L.; Al Haj Zen, A.; Fourches, D.; Muratov, E.; Tropsha, A.; Morris, J.; Teicher, B. A.; Kunkel, M.; Polley, E.; Lackey, K. E.; Atkinson, F. L.; Overington, J. P.; Bamborough, P.; Müller, S.; Price, D. J.; Willson, T. M.; Drewry, D. H.; Knapp, S.; Zuercher, W. J. Comprehensive Characterization of the Published Kinase Inhibitor Set. *Nat Biotechnol* **2016**, *34* (1), 95–103. https://doi.org/10.1038/nbt.3374.

(7) Vasta, J. D.; Corona, C. R.; Wilkinson, J.; Zimprich, C. A.; Hartnett, J. R.; Ingold, M. R.; Zimmerman, K.; Machleidt, T.; Kirkland, T. A.; Huwiler, K. G.; Ohana, R. F.; Slater, M.; Otto, P.; Cong, M.; Wells, C. I.; Berger, B.-T.; Hanke, T.; Glas, C.; Ding, K.; Drewry, D. H.; Huber, K. V. M.; Willson, T. M.; Knapp, S.; Müller, S.; Meisenheimer, P. L.; Fan, F.; Wood, K. V.; Robers, M. B. Quantitative, Wide-Spectrum Kinase Profiling in Live Cells for Assessing the Effect of Cellular ATP on Target Engagement. *Cell Chem Biol* **2018**, *25* (2), 206-214.e11. https://doi.org/10.1016/j.chembiol.2017.10.010.

**7-Hydroxy-MN**

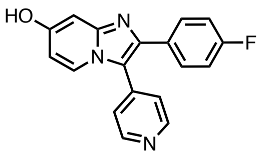

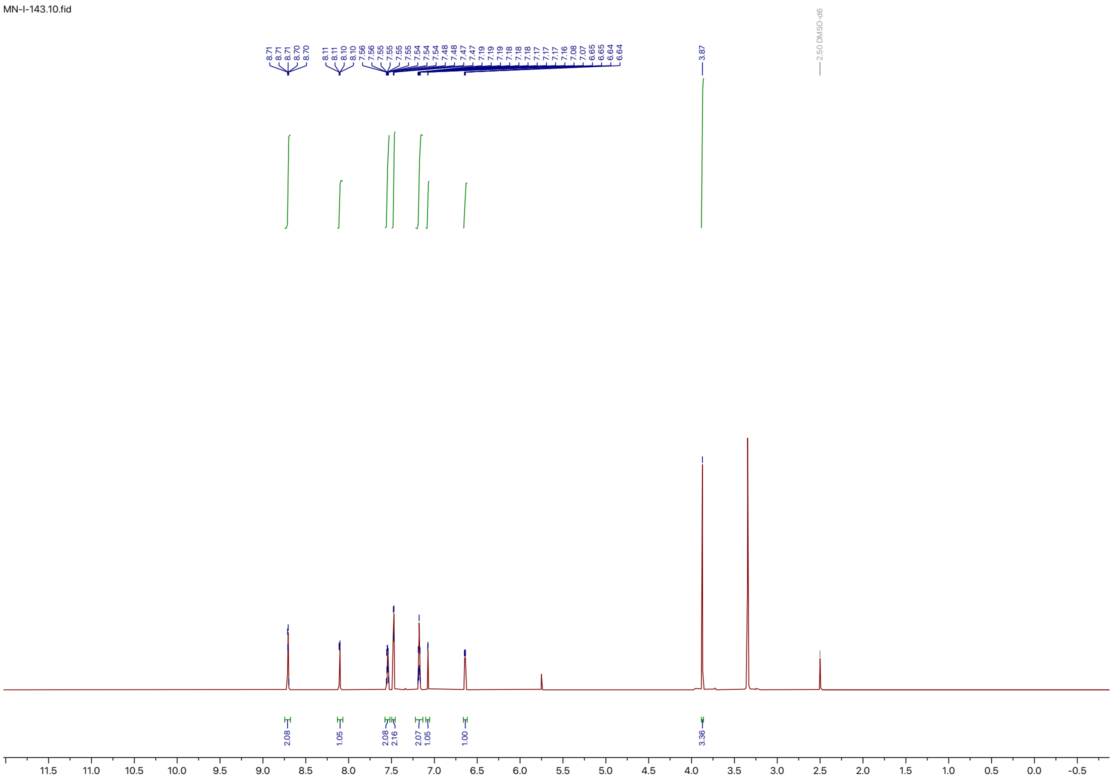
MN-I-143.10.fid
8.71
8.71
8.71
8.70
8.70
8.11
8.11
8.10
8.10
7.56
7.56
7.55
7.55
7.55
7.55
7.54
7.54
7.54
7.48
7.48
7.47
7.47
7.19
7.19
7.19
7.18
7.18
7.18
7.17
7.17
7.17
7.16
7.08
7.07
6.65
6.65
6.64
6.64
3.87
2.50 DMSO-d6
2.08
1.05
2.08
2.16
2.07
1.05
1.00
3.36
11.5
11.0
10.5
10.0
9.5
9.0
8.5
8.0
7.5
7.0
6.5
6.0
5.5
5.0
4.5
4.0
3.5
3.0
2.5
2.0
1.5
1.0
0.5
0.0
-0.5

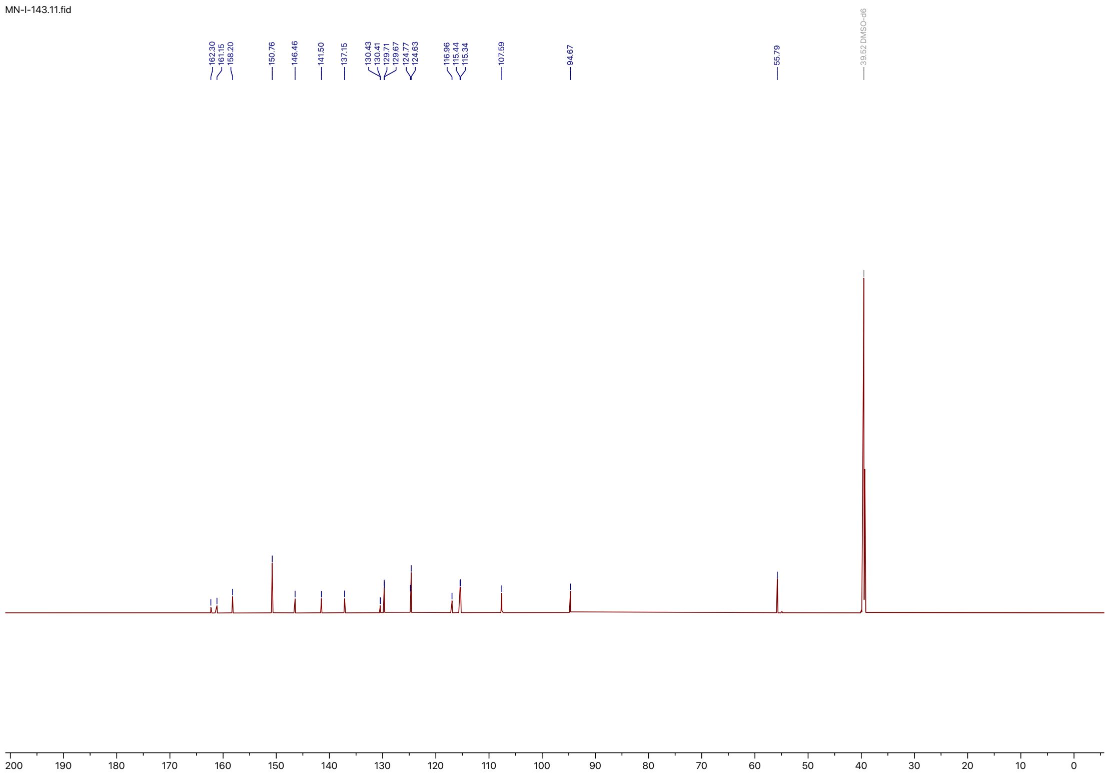
MN-I-143.11.fid
162.30
161.15
158.20
150.76
146.46
141.50
137.15
130.43
130.41
129.71
129.67
124.77
124.63
116.96
115.44
115.34
107.59
94.67
55.79
39.52 DMSO-d6
200 190 180 170 160 150 140 130 120 110 100 90 80 70 60 50 40 30 20 10 0

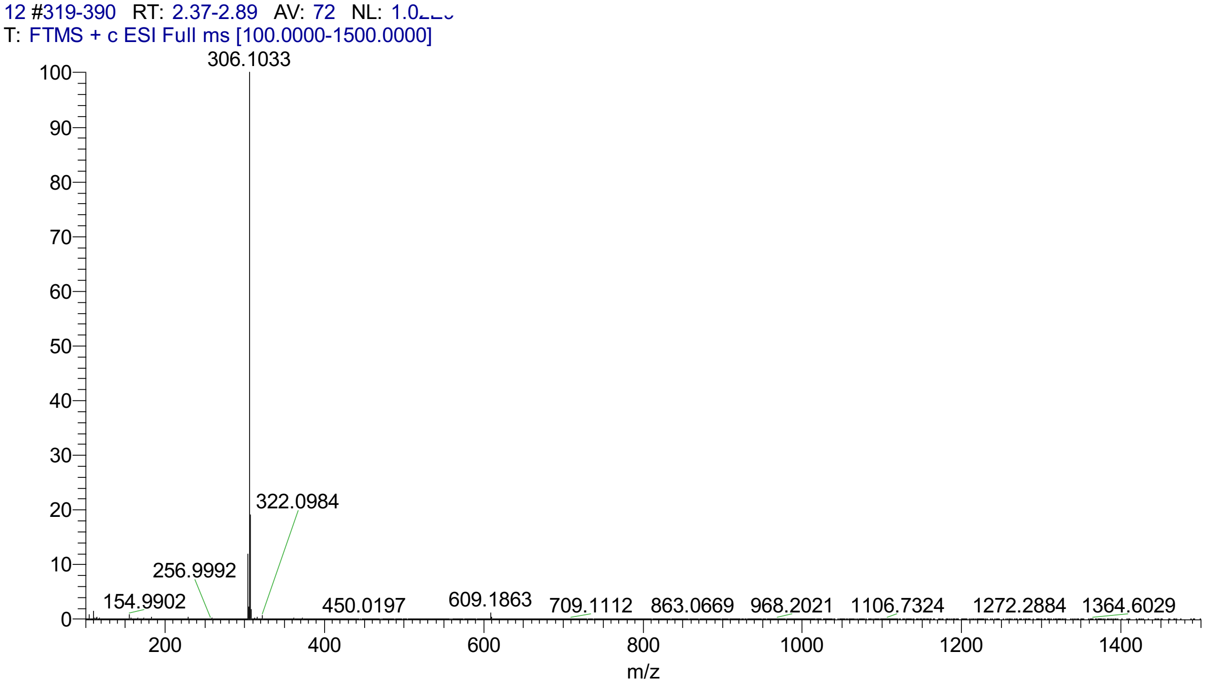
12 #319-390 RT: 2.37-2.89 AV: 72 NL: 1.0
T: FTMS + c ESI Full ms [100.0000-1500.0000]
306.1033
322.0984
256.9992
154.9902
450.0197
609.1863
709.1112
863.0669
968.2021
1106.7324
1272.2884
1364.6029
m/z

**7-(2-Aminoethoxy)-MN**

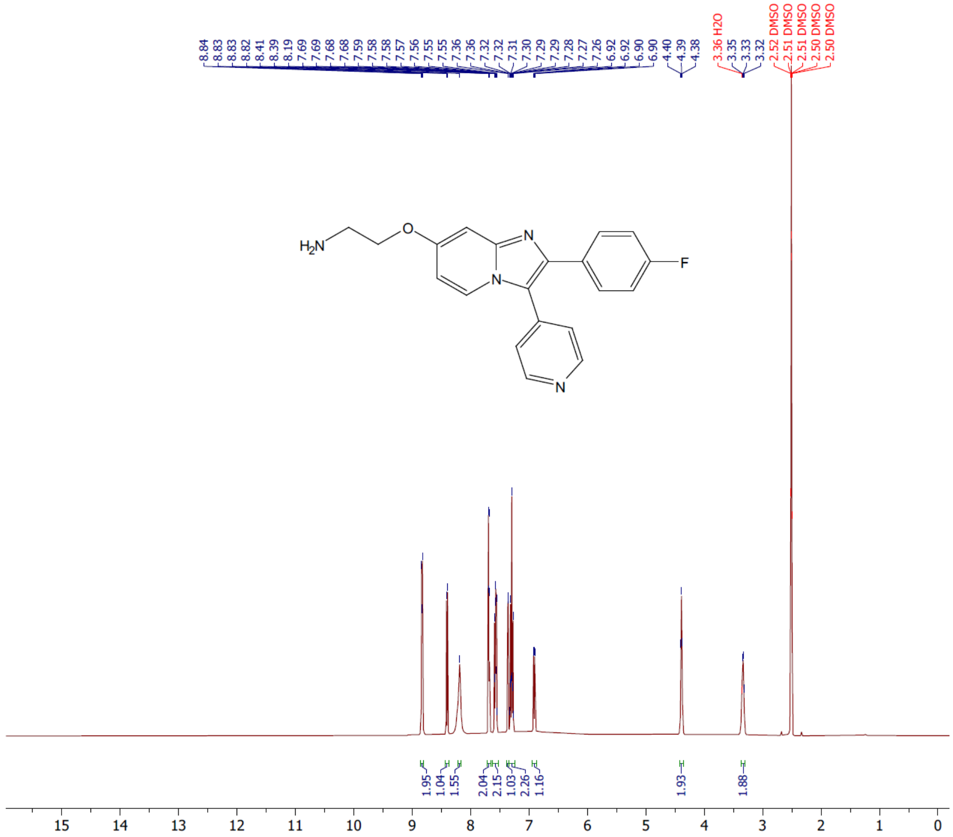


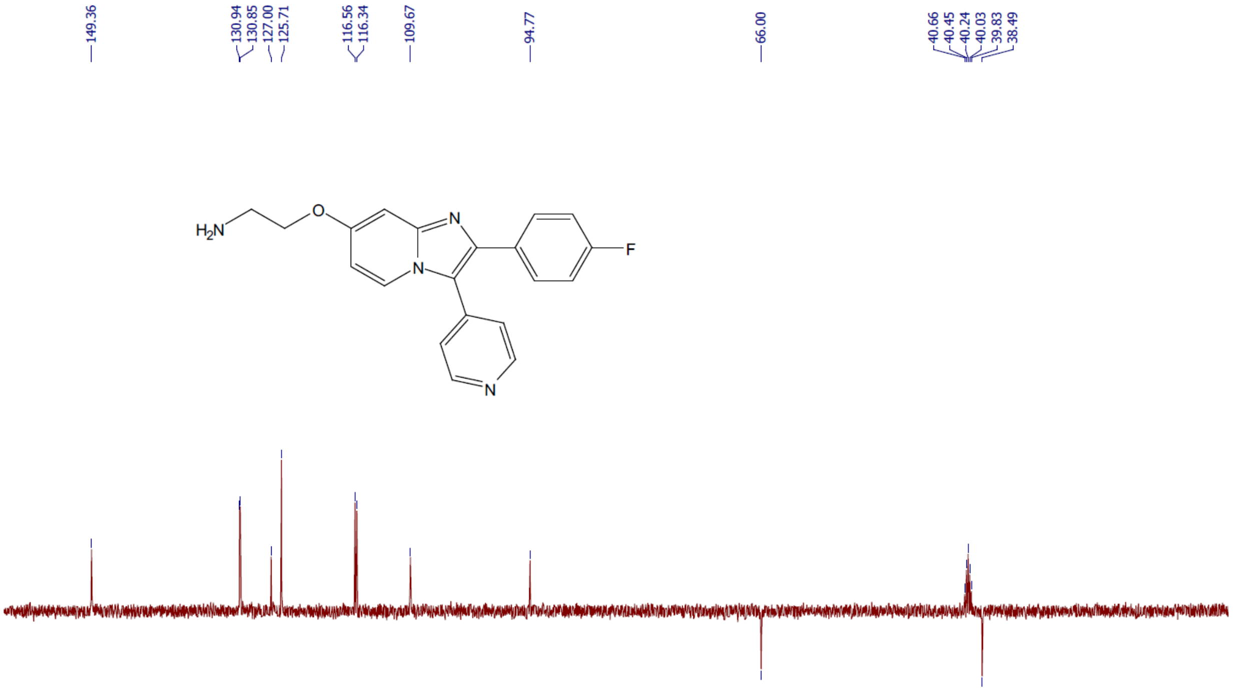

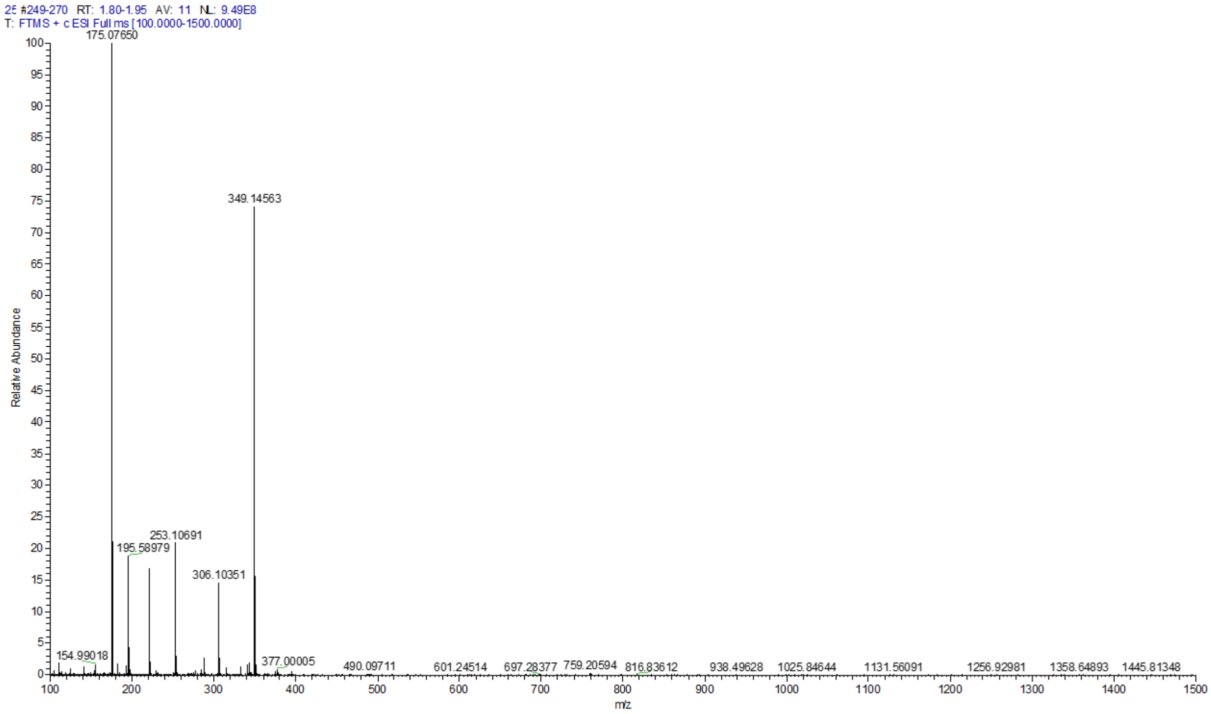

25 #249-270 RT: 1.80-1.95 AV: 11 NL: 9.49E8
T: FTMS + c ESI Full ms [100.0000-1500.0000]
175.07650
349.14563
253.10691
195.58979
306.10351
154.99018
377.00005
490.09711
601.24514
697.28377
759.20594
816.83612
938.49628
1025.84644
1131.56091
1256.92981
1358.64893
1445.81348
Relative Abundance
m/z

**Boc-amino-PEG-MN**

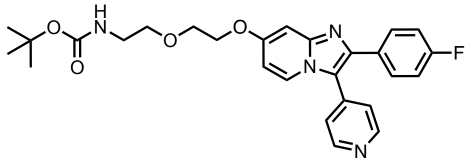

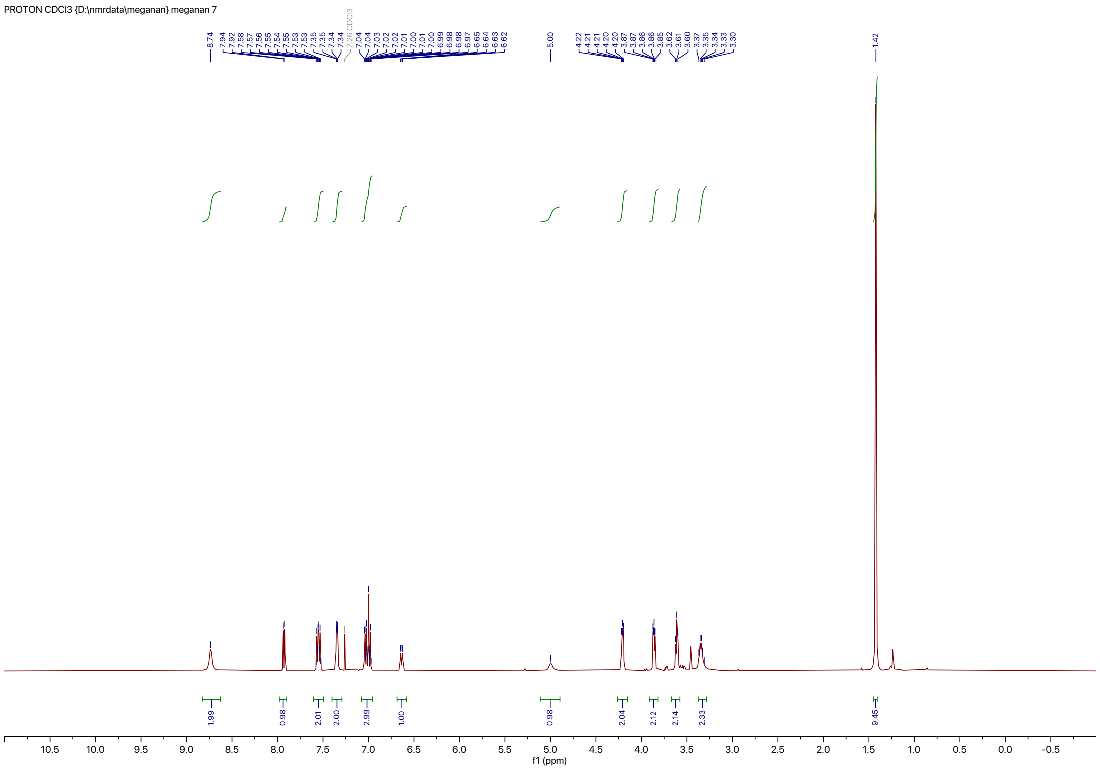

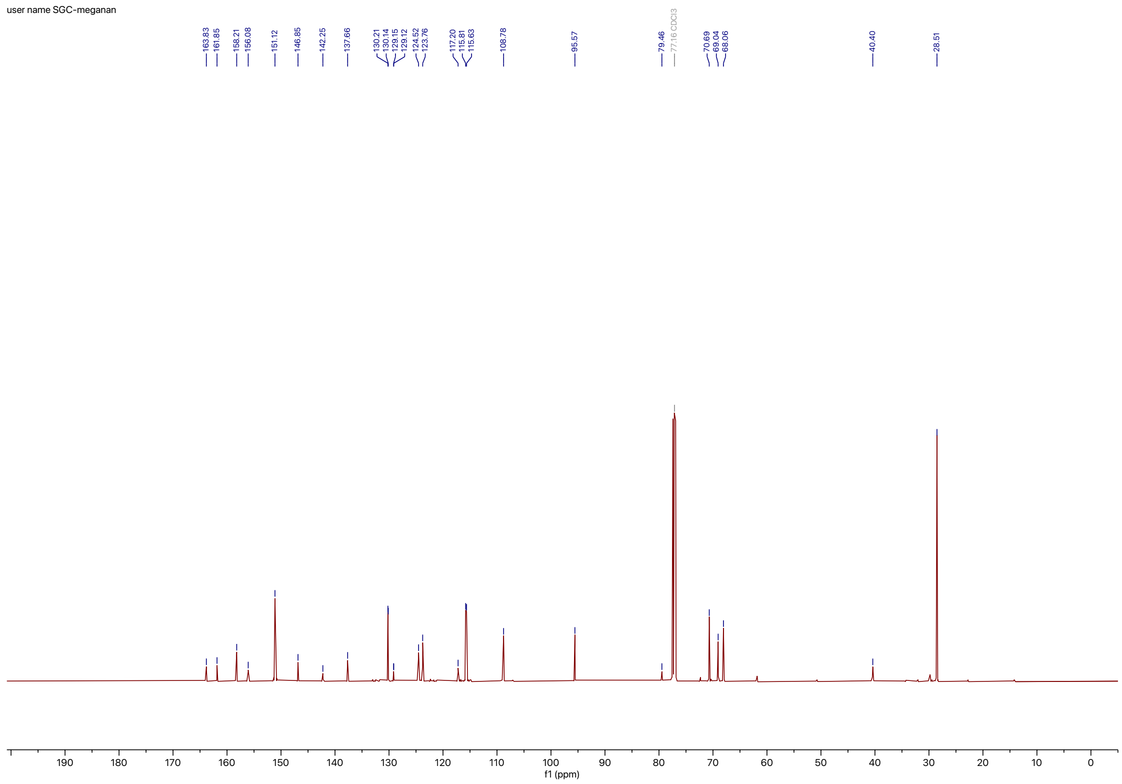
user name SGC-meganan
163.83
161.85
158.21
156.08
151.12
146.85
142.25
137.66
130.21
130.14
129.15
129.12
124.52
123.76
117.20
115.81
115.63
108.78
95.57
79.46
77.16 CDCl3
70.69
69.04
68.06
40.40
28.51
f1 (ppm)

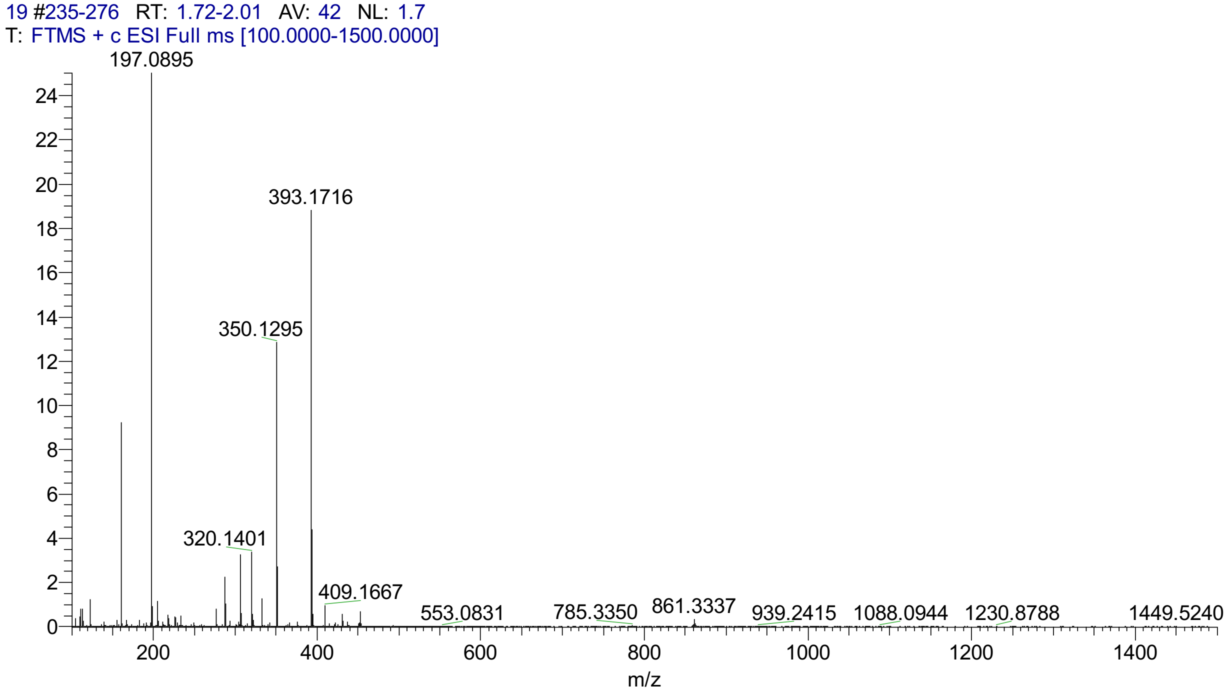
19 #235-276 RT: 1.72-2.01 AV: 42 NL: 1.7
T: FTMS + c ESI Full ms [100.0000-1500.0000]
197.0895
393.1716
350.1295
320.1401
409.1667
553.0831
785.3350
861.3337
939.2415
1088.0944
1230.8788
1449.5240
m/z

### MN-Bodipy

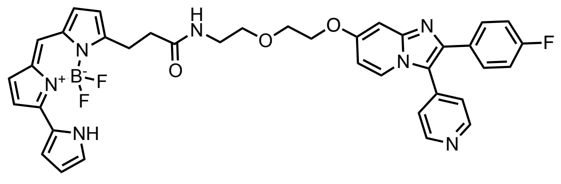


user name SGC-meganan

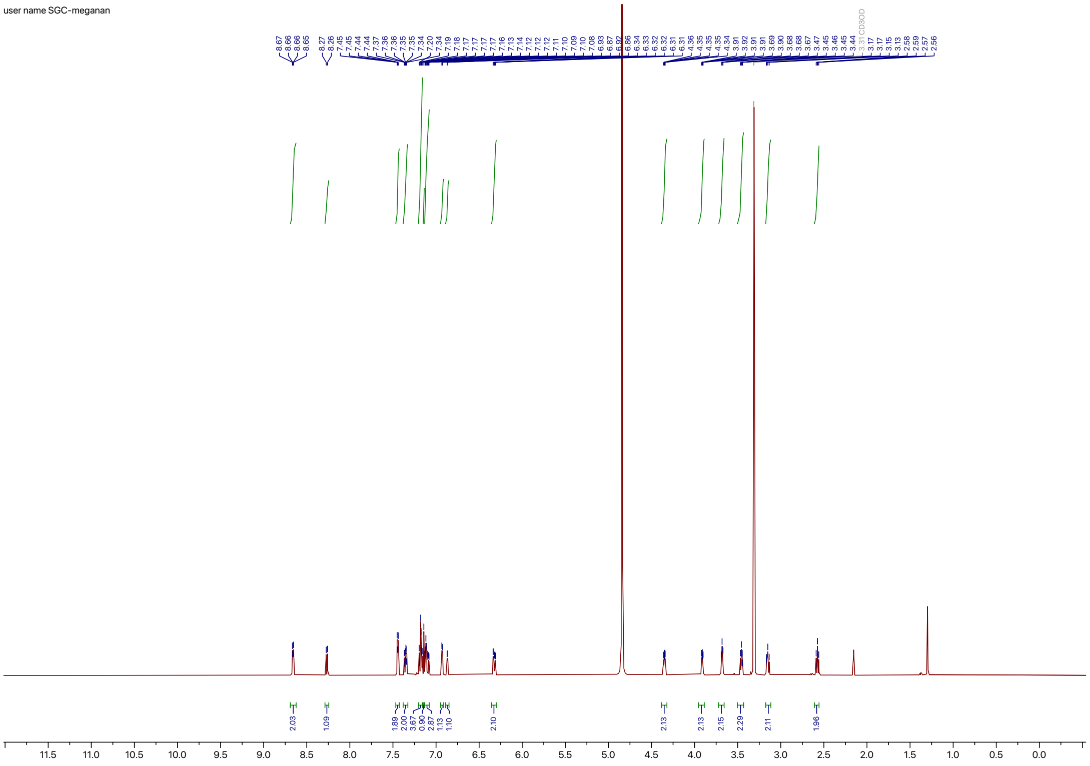

user name SGC-meganan

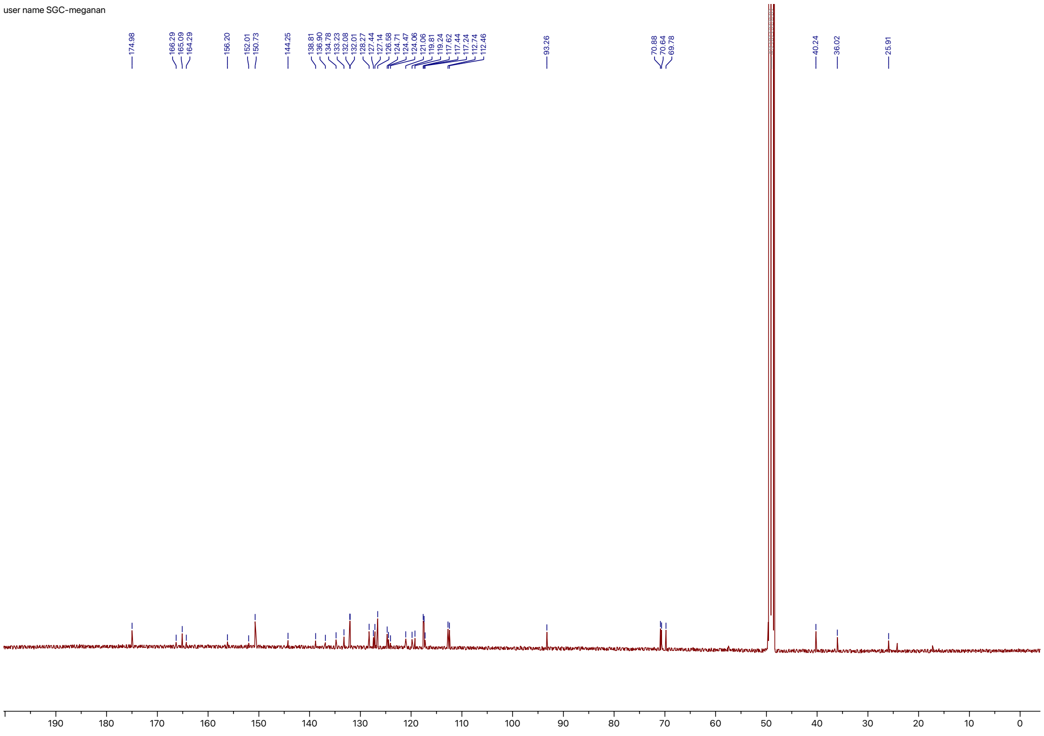

user name SGC-meganan

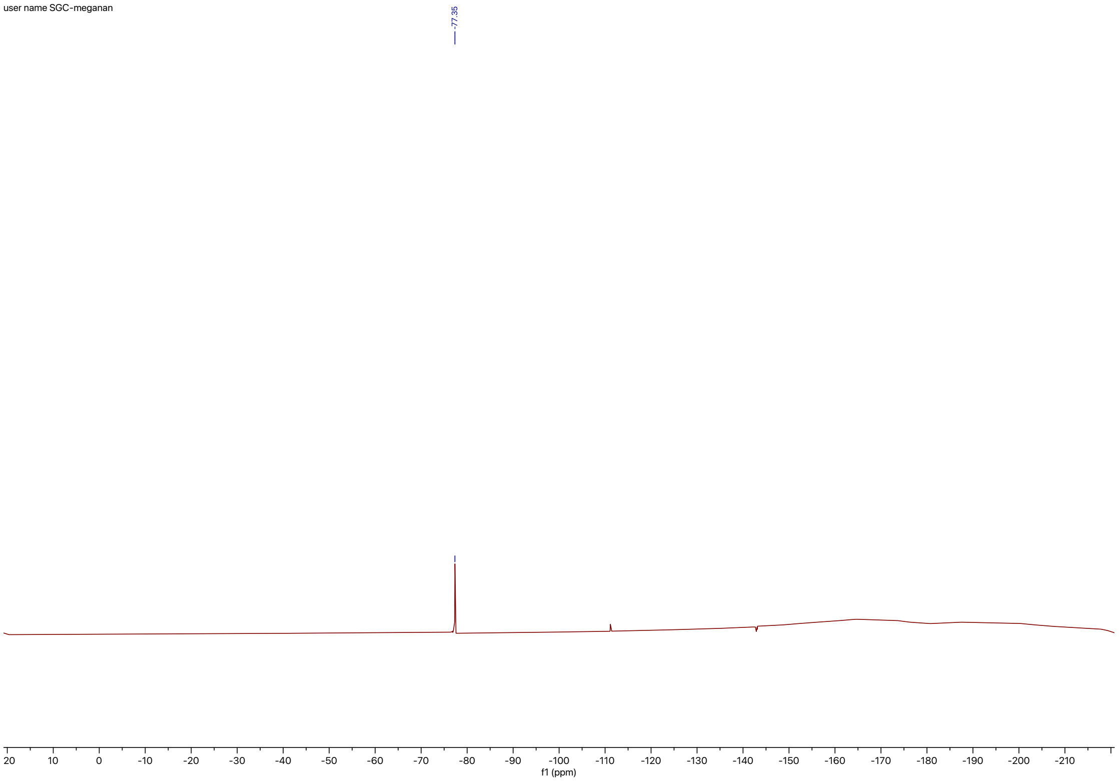

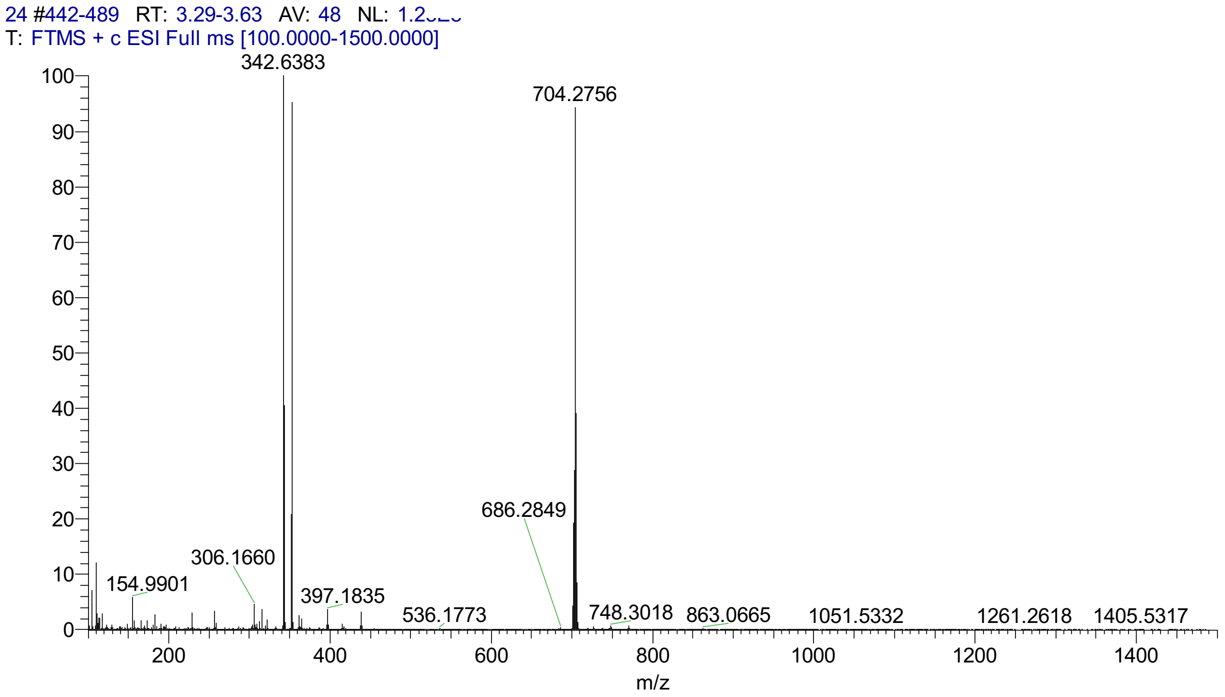
24 #442-489 RT: 3.29-3.63 AV: 48 NL: 1.2
T: FTMS + c ESI Full ms [100.0000-1500.0000]
342.6383
704.2756
686.2849
306.1660
154.9901
397.1835
536.1773
748.3018
863.0665
1051.5332
1261.2618
1405.5317
m/z
